# Variation in mutation, recombination, and transposition rates in *Drosophila melanogaster* and *Drosophila simulans*

**DOI:** 10.1101/2022.09.12.507595

**Authors:** Yiguan Wang, Paul McNeil, Rashidatu Abdulazeez, Marta Pascual, Susan E. Johnston, Peter D. Keightley, Darren J. Obbard

## Abstract

Mutation, recombination, and transposition occurring during meiosis provide the variation on which natural selection can act and the rates at which they occur are important parameters in models of evolution. The *de novo* mutation rate determines levels of genetic diversity, responses to ongoing selection, and levels of genetic load. Recombination breaks up haplotypes and reduces the effects of linkage, helping to spread beneficial alleles and purge deleterious ones. Transposable elements (TE) selfishly replicate themselves through the genome, imposing fitness costs on the host and introducing complex mutations that can affect gene expression and give rise to new genes. However, even for key evolutionary models such as *Drosophila melanogaster* and *D. simulans* few estimates of these parameters are available, and we have little idea of how rates vary between individuals, sexes, populations, or species. Here, we provide direct estimates of mutation, recombination, and transposition rates and their variation in a West African and a European population of *D. melanogaster* and a European population of *D. simulans*. Across 89 flies, we observe 58 single nucleotide mutations, 286 crossovers, and 89 TE insertions. Compared to the European *D. melanogaster*, we find the West African population has a lower mutation rate (1.67 *vs*. 4.86 × 10^−9^ site^−1^ gen^−1^) and transposition rate (8.99 *vs*. 23.36 × 10^−5^ copy^−1^ gen^−1^), but a higher recombination rate (3.44 *vs*. 2.06 cM/Mb). The European *D. simulans* population has a similar mutation rate to European *D. melanogaster* but a significantly higher recombination rate and a lower but not significantly different transposition rate. Overall, we find paternal-derived mutations are more frequent than maternal ones in both species.

**Highlights:**

- *De novo* mutation rates are 1.67 × 10^−9^ site^−1^ gen^−1^ (95% HPD CI: 0.54 – 3.14 × 10^−9^), 4.86 × 10^−9^ site^−1^ gen^−1^ (2.11 – 8.02 × 10^−9^), and 4.51 × 10^−9^ site^−1^ gen^−1^ (1.94 – 7.75 × 10^−9^) for the West African *D. melanogaster*, the European *D. melanogaster* and the European *D. simulans* population, respectively.
- In females, recombination rates in the absence of large genomic inversions are 3.44 cM/Mb (2.72 – 4.18), 2.06 cM/Mb (1.57 - 2.57), and 3.04 cM/Mb (2.45 - 3.73) for the three populations, respectively. There was no strong evidence of recombination observed in males.
- Mutations (SNMs and indels) are male-biased.
- The West African *D. melanogaster* population has a lower TE activity than the other populations and *CMC-Transib* is the dominant active TE. The European *D. melanogaster* population has multiple active TEs: *Gypsy, CMC-Transib, Pao, Jockey* and *hAT-hobo*; while in European *D. simulans*, they are *Gypsy, CMC-Transib, Pao, hAT-hobo, Copia* and *TcMar-Mariner*.

## 1. Introduction

Mutation is the ultimate source of all genetic variation, and the mutation rate consequently plays a key role in evolutionary processes (Lynch 2010; Lynch, et al. 2016). Although the germline mutation rate is thought to be lower than the somatic mutation rate (Milholland, et al. 2017), only germline mutations can be inherited, and so have greater importance in population and evolutionary genetics. It has been hypothesised that the germline mutation rate in multicellular eukaryotes may be explained by an equilibrium between natural selection to minimise it and the power of genetic drift to overcome the effect of selection (Lynch, et al. 2016). However, to date, there are surprisingly few estimates of germline mutation rate from multicellular eukaryotes, and those estimates that we do have are patchily distributed across the tree of life (reviewed by Yoder and Tiley 2021). On the one hand, mutation rates appear to differ markedly between some distantly related species. For example, primate mutation rates are generally greater than 10 × 10^−9^ site^−1^ gen^−1^ (Rahbari, et al. 2016; Jonsson, et al. 2017; Lindsay, et al. 2019), whereas insect mutation rates are generally lower than 6 × 10^−9^ site^−1^ gen^−1^ (Keightley, et al. 2014; Keightley, et al. 2015; Yang, et al. 2015; Liu, et al. 2017). On the other hand, some very distantly-related lineages, such as *Drosophila* and *Heliconius* that are separated by ~290 million years of evolution (Misof, et al. 2014; Suvorov, et al. 2022), have estimates that are not significantly different from each other (Keightley, et al. 2014; Keightley, et al. 2015). The apparent variation in mutation rate may, in part, reflect variation within species combined with very limited sampling. For example, in humans—for which many direct estimates are available—estimates range from 9.6 × 10^−9^ (Campbell, et al. 2012) to 21.7 × 10^−9^ site^−1^ gen^−1^ (O’Roak, et al. 2012) and parental age at conception is positively correlated with the number of mutations in the offspring (Kong, et al. 2012; Kaplanis, et al. 2021). For *Drosophila melanogaster*, there are relatively few estimates, ranging from 2.8 × 10^−9^ site^−1^ gen^−1^ to 6.0 × 10^−9^ site^−1^ gen^−1^ (Haag-Liautard, et al. 2007; Keightley, et al. 2009; Schrider, et al. 2013; Keightley, et al. 2014; Huang, et al. 2016; Sharp and Agrawal 2016; Assaf, et al. 2017), but there has been no formal attempt to quantify variation in mutation rate between the sexes or among wild individuals or populations.

Apparent variation in mutation rate may also be partly attributable to a lack of precision in the estimates, as the challenge of accurately estimating the mutation rate can be formidable when one or fewer *de novo* mutations is expected per offspring and when there are millions of segregating sites— as in *D. melanogaster* (Keightley, et al. 2014). As a consequence of this, most studies in model species have used a ‘mutation accumulation’ (MA) approach, in which selection is relaxed for multiple generations in fully inbred sib-mated lines and multiple mutations are counted at the end of the experiment (Sung, et al. 2015; Uchimura, et al. 2015; Oppold and Pfenninger 2017). Unfortunately this approach cannot detect recessive lethal mutations and may conceivably be biased by the accumulation of deleterious variants that themselves affect mutation rates (Baer, et al. 2007). In contrast, pedigree or parent-offspring trio studies, such as those as widely used for humans and other large mammals, give a direct and relatively unbiased estimate of the mutation rate (Segurel, et al. 2014). This approach has rarely been applied in *Drosophila* (but see Keightley, et al. 2014; Krasovec 2021), although it provides the opportunity to obtain less biased estimates of the mutation rate and how it varies within and between species.

Recombination also plays a central role in creating variation, by breaking linkage between segregating mutations and facilitating their independent fixation or loss (Webster and Hurst 2012). However, in contrast to mutation rates, recombination rates have frequently been estimated in mapping studies. These have shown that recombination rates vary within and between chromosomes, individuals, sexes, populations, and species, and the variation can be attributed to both genetic factors (e.g. rate and landscape modifier loci, genome architecture and chromatin structure) and environmental factors (e.g. age, temperature and pathogen infection) (Stapley, et al. 2017). Genome wide estimates of recombination rate are available for several *Drosophila* species, including a *D. serrata* estimate of *ca*. 1.4 cM/Mb (Stocker, et al. 2012), *D. pseudoobscura* of 3.3 - 4.6 cM/Mb, *D. miranda* of 4.9 – 6.1 cM/Mb (Heil, et al. 2015), *D. persimilis* of 4.1 cM/Mb for autosomes and 5.0 cM/Mb for X chromosome (Stevison and Noor 2010), *D. virilis* of 4.6 cM/Mb (Hemmer, et al. 2020) and *D. melanogaster* of 2.5 cM/Mb (Comeron, et al. 2012). However, experimental studies of *Drosophila* have rarely looked at variation in genome-wide recombination rates in natural populations of outbred individuals, and wild populations may differ systematically from populations that have been strongly selected by adaptation to the laboratory (Aggarwal, et al. 2021).

Transposition by mobile genetic elements (transposable elements; TEs), while perhaps less widely considered by evolutionary studies as an important source of variation, can occur at a rate similar to that of the nucleotide mutation rate (Adrion, et al. 2017). New TE insertions can be major source of loss-of-function mutations (Hirsch and Springer 2017), including those that directly disrupt genes and those that indirectly lead to ectopic chromatic formation (Wells and Feschotte 2020), and over evolutionary time TEs have surprisingly often been ‘domesticated’ into important roles in the host genome (Chuong, et al. 2017). Using *in situ* hybridization in *D. melanogaster*, some early studies revealed that the transposition rate varied by several order of magnitude, ranging from 10^−5^ copy^−1^ gen^−1^ to 10^−2^ copy^−1^ gen^−1^ (Nuzhdin and Mackay 1995; Pasyukova, et al. 1998; Maside, et al. 2000; Diaz-Gonzalez, et al. 2011). This variation is partly due to differences in activity between TE families, for example, *INE-1* has been inactive for millions of years in *D. melanogaster*, whereas *Transib* is recently active (Kapitonov and Jurka 2003). TE activity also varies among populations or chromosomes (Adrion, et al. 2017), and is affected by factors such as stress and aging (Capy, et al. 2000; Guerreiro 2012; Chuong, et al. 2017; Horvath, et al. 2017). It has also been claimed that colonization of new habitats could induce transposition in some *Drosophila* species, for example in *D. buzzatii* and *D. subobscura* (Guerreiro, et al. 2008; Guerreiro and Fontdevila 2011), highlighting the potential influence of environmental stress and emphasising differences in TE activity between laboratory and wild conditions.

Here, we directly estimate the *de novo* mutation rate, recombination rate, and rate of TE insertion in low-complexity regions by sequencing parents and offspring from 18 families of full-sib *D. melanogaster* and *D. simulans*. By sequencing unrelated parents that were either wild collected themselves (the majority of the fathers) or the offspring of wild-collected flies (mothers), along with their first-generation offspring, our estimates are less subject than many previous studies to the impacts of selection, inbreeding, homozygosity, and the accumulation of deleterious mutations. We find a higher mutation rate but lower recombination rate in the European population of *D. melanogaster* than in their West African counterpart. The European *D. simulans* population has a similar mutation rate, but a higher recombination rate, than the European *D. melanogaster*. TE activity differs dramatically among the three populations, the European *D. melanogaster* exhibiting a higher transposition rate than the West African population. These direct estimates of mutation, recombination, and transposition rates in fully wild fly genomes not only highlight the variation in rates among populations and individuals, but also provide improved parameter estimates for future evolutionary studies.

## 2. Materials and Methods

### 2.1 Fly Crosses and sequencing

West African *D. melanogaster* were netted by RA from fermented banana bait that had been exposed for 48 hours in Samaru market in Zaria, northern Nigeria (28^th^ September 2020; 11.1611 N, 7.6471 E). European *D. melanogaster* were netted by DJO from yeasted banana bait that had been exposed for around 5 days in a rural garden in Sussex, southeast England (4^th^ to 10^th^ August 2020; 51.0998 N, 0.1644 E). European *D. simulans* were netted by MP from decomposing fallen fruit in an organic apple and pear orchard in Gimenells, Northern Spain (21^st^ August 2020; 41.6564 N, 0.3885 E). In each case the landowner’s permission was obtained before collection, and this work was approved by the University of Edinburgh School of Biological Science ethics committee.

For the West African flies, we crossed the virgin male and female offspring from different wild-collected (i.e. wild-mated) females with each other. For the European flies, we crossed virgin female offspring of wild-collected females with wild-collected males. All ‘parental’ flies for sequencing were therefore unrelated and genetically wild, i.e. no more inbred or related than naturally occurring flies. All progeny groups comprised outbred full-sibships. Once sufficient numbers of F1 offspring were available, we froze the parents at −80°C until DNA extraction. All flies were maintained at 16:8 light:dark cycle at 20°C on a modified ‘Lewis’ medium. In total, we sequenced six families comprising two parents and five F1 offspring (males) from each of the three populations (West African and European *D. melanogaster*, European *D. simulans*), giving 125 flies in total (36 parents and 89 offspring – one offspring individual failed sequencing).

DNA was extracted from single flies using Qiagen Blood and Tissue kits (Qiagen Ltd., UK) according to the manufacturer’s protocol, and provided to the Centre for Genomic Research at the University of Liverpool (UK) for library preparation and sequencing. Libraries were prepared using NEBNext Ultra II FS Kits (New England Biolabs (UK) Ltd) with an insert size of approximately 350bp. Paired-end (2 × 150bp) sequencing was performed using Illumina NovaSeq with S4 chemistry, and raw fastq files were trimmed for adapter sequences using Cutadapt-version 1.2.1 (Martin 2011). Reads were further trimmed for quality using Sickle (https://github.com/najoshi/sickle, version1.200), with a minimum window quality score of 20. After trimming, we retained only paired reads, giving an average of 38.3 million pairs for each sample, with a range from 20.0 to 81.1 million. The median coverages were all above 30X (except one with 28X) and the X chromosome coverages in males were half of that in autosomes.

### 2.2 Read mapping and variant calling

We used BWA-MEM (version 0.7.17-r1188; Li 2013) to map the trimmed reads to the reference genomes of *D. melanogaster* (version r6.42, flybase.org) and *D. simulans* (version GCF_016746395.2_Prin_Dsim_3.1, NCBI) using default parameters. We used Picard MarkDuplicates (https://github.com/broadinstitute/picard, version 2.26.2) to identify and remove duplicate reads in the BAM files with parameters ‘REMOVE_DUPLICATES=true ASSUME_SORTED=true VALIDATION_STRINGENCY=LENIENT’. We used BCFtools-version 1.11 (Li 2011) to call variants for all the samples in each family with parameters ‘-a DP,AD,ADF,ADR --max-depth 250 --min-MQ 20’. Given the haploid state of the X and Y chromosomes in male samples, we configured a ploidy file using ‘--ploidy-file’ in BCFtools. Different chromosome arms were called separately and then concatenated using ‘bcftools concat’. We used GCTA (version 1.93.2; Yang, et al. 2011) to examine the relatedness of the parental flies, which calculates genetic relatedness matrix (GRM) between pairs of individuals *j* and *k*:

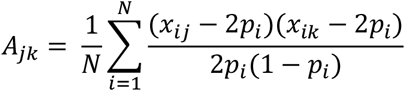

where *N* is the total number of SNPs, *x_ij_* is the copy of reference allele for the *i^th^* SNP in individual *j*, and *p_i_* is the population allele frequency for the *i^th^* SNP. We also used seqinr (version 4.2-5; Charif and Lobry 2007) to estimate the genome-wide synonymous nucleotide diversity from the inferred parental haplotypes (*π_s_*).

### 2.3 Filtering of candidate single-nucleotide and short indel mutations

The central challenge in calling rare heterozygous *de novo* mutations in highly diverse diploid genomes is to distinguish true mutations from read-mapping errors (Keightley, et al. 2014; Bergeron, et al. 2022). For each family, we used GATK-version 4.2.2 (McKenna, et al. 2010) ‘SelectVariants’ to select single nucleotide variants (SNVs) in the VCF file and then used ‘VariantsToTable’ to extract relevant information for each SNV, including chromosomes (CHROM), positions (POS), reference allele (REF), alternative allele (ALT), genotype (GT), depth (DP), genotype quality (GQ), allele depth (AD), allele depth on forward reads (ADF) and allele depth on reverse reads (ADR). We then filtered the SNVs using the criteria in **BOX 1A** to identify sites of candidate *de novo* mutations.

#### BOX 1. Filtering strategies for SNMs (A) and indels (B)

**A**: To filter SNMs, we applied the following criteria:

For both parents we required that:

- The site was called homozygous for the reference allele (GT)
- The read depth (DP) was in the range 10X – 150X
- The genotype quality (GQ) was ≥ 50
- The site was ‘pure’, i.e. no reads supporting the candidate mutation (AD = 0)

For each focal offspring we required that:

- The site was called heterozygous (GT)
- The read depth (DP) was in the range 10X – 150X
- The genotype quality (GQ) was ≥ 50
- The proportion of reads supporting the alternative allele relative to the total depth at this site (allelic balance, AB) was in the range 0.2 – 0.8
- The number of reads supporting the candidate mutation was ≥ 5 (AD ≥ 5)
- The candidate mutation was supported by both forward (ADF > 0) and reverse (ADR > 0) reads

For non-focal offspring in the family, we required that:

- The site was called homozygous for the reference allele (GT)
- No offspring had more than one read supporting the candidate mutation
- No more than two offspring had any reads supporting the candidate mutation

For all other families in the same population, we required that:

- No individual was called heterozygous (GT)

**B:** To filter indels, we applied the following criteria:

For both parents we required that:

- The number of reads covering the indel location (NR) was between 10 - 150
- No read in either parent contained the candidate indel

For the focal offspring we required that:

- The number of reads covering the indel location (NR) was between 10 - 150
- The number of reads containing the indel (NV) was ≥ 5
- The allelic balance was between 0.15 - 0.85

For non-focal offspring:

- No read in any offspring contained the candidate indel

For the other families in the same population:

- No read in any individual contained the candidate indel

We additionally tailored the filtering approach for SNVs on X and Y chromosomes in males, as follows. For the X chromosome, we did not filter the father except to require that the site was ‘pure’ (i.e. no reads supporting the candidate mutation; AD = 0), the allelic balance (AB) for the focal (male) offspring was 1, and the read depth was 5X - 75X (i.e. half the range for diploid filtering). For the Y chromosome, we required that the father was ‘pure’, with read depth 5X – 75X and that the focal offspring was AB = 1.

To filter candidate short indel mutations (those less than 50 bp), we re-called indels for each of the families using platypus-version 0.8.1.2 (Rimmer, et al. 2014) with parameters ‘--genSNPs=0 -- countOnlyExactIndelMatches=1’. Then we used VCFtools-version 0.1.16 (Danecek, et al. 2011) to extract biallelic indels from the VCF files, and used GATK ‘VariantsToTable’ to extract chromosomes (CHROM), positions (POS), reference allele (REF), alternative allele (ALT), the number of reads covering variant location (NR), and the number of reads containing the variant in this sample (NV). We then used the criteria in **BOX 1B** to identify sites of candidate *de novo* short indels.

Similar to the filtering for SNVs above, we halved the required number of reads (NR) to filter candidate variants on X and Y chromosomes in male offspring, and we required that the allelic balance was over 0.5. We did not filter the number of reads (NR) in the father, but we did require NV to be 0 if the candidate variant were on the X chromosome (i.e. the sample was ‘pure’).

Finally, for each family, we manually examined all of the candidate SNMs and short indel mutations using the Integrated Genomics Viewer (IGV; Robinson, et al. 2011). To be accepted, a candidate mutation was required to fulfil the following criteria. First, we ensured the candidate mutation was absent from all parental reads, and satisfied the criteria described above in the non-focal offspring. Second, the candidate mutation and its surrounding genomic markers (such as SNP or indel alleles, if present) had to be phased into only two haplotypes if on autosomes, or one haplotype if on sex chromosomes in a male. This step also allowed us to trace the parent of origin for the majority of mutations. Third, if a read containing the candidate mutation was present in any non-focal offspring, and it carried some other genomic markers, then the association between the candidate mutation and these genomic markers should not be identified in the focal offspring (Keightley, et al. 2014). Last, although we didn’t exclude putative mutations in the close vicinity of indels in the automated pipeline outlined above, we did verify these manually, and candidates were excluded if they could be perfectly resolved by moving bases around the indels (Keightley, et al. 2014). All scripts are available via GitHub:https://github.com/Yiguan/muDmelDsim.

### 2.4 Simulation and defining ‘callable’ sites

Unevenness in mapped read depth, for example due to variation in ‘mappability’, means that some fraction of the genome will not be accessible to mutation detection. Most estimates of mutation rate account for this missing fraction by using only the ‘callable’ sites as the denominator when calculating per-site rates (i.e. the sites meeting the depth and quality threshold in all relevant individuals). However, the complex filtering required to achieve high specificity (**Box 1**) can reduce detection sensitivity in other ways that may be harder to capture, in part because they are contingent on the mutations that arise. For example, as mismatches can reduce the probability that a read maps successfully, especially in a high-diversity genome, the mapped read depth may itself depend on the presence of new mutations. Even more subtly, a ‘purity’ requirement means that a site called as reference homozygous AA, but which has a very small fraction of reads supporting a C allele, is ‘callable’ if the *de novo* mutation is A→G, but not if it is A→C. One solution to this has been to simulate ‘synthetic mutations’ in the empirical reads, and re-run the mapping and mutation-detection pipeline to estimate its overall sensitivity (Keightley, et al. 2014). This sensitivity estimate can then be used to ‘correct’ the overall rate estimate. Here we take this approach one step further, and use these simulations to directly define the proportion of the genome that is ‘callable’, simply as the proportion of synthetic mutations that can be re-called by the whole pipeline.

To simulate SNMs, we randomly selected 100,000 ‘synthetic mutation’ sites across the combined genomes of the five offspring in each family, ensuring that sites were at least 500bp apart and setting the number of sites per chromosome in proportion to the chromosome length. We then extracted all of the reads covering each of these sites and edited the bases in these reads at each chosen site to create a synthetic mutation. To mimic the number of non-reference alleles and their linkage to true standing variation, we followed two rules. First, if there were SNP markers linked the chosen site that could be used to phase the reads, we randomly selected one parental haplotype and edited all reads belonging to this haplotype. Second, if there were no available SNP markers on the target reads, we randomly selected the number of reads to be edited based on the empirical distribution of read frequencies for true heterozygous sites, given the read depth. We then replaced the corresponding reads in the original BAM files with the reads that had been edited and used ‘samtools fastq’ to convert the edited BAM files to paired-end read files. Together with the original read files for the parents in each family we then re-ran our complete pipeline, including mapping, variant calling and filtering. If a synthetic mutation was successfully recovered by the full pipeline, then the synthetic mutation site was deemed callable. We then estimated the *total* callable sites as:

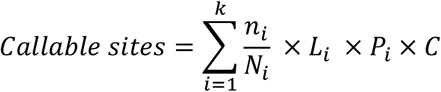

where, *k* is the number of chromosomes; *n_i_* is the number of callable sites in a total of *N_i_* sites we simulated on the *ith* chromosome; *L_i_* is the length of the reference genome for the *ith* chromosome; *C* is the number of offspring in the family; *P_i_* is the number of ploidy for the *ith* chromosome in each offspring.

We also simulated synthetic short indels in our data, analogous to our approach for the simulation of SNMs. However, given the limits of short-read data, we limited the length of the synthetic indels to a maximum of 25bp. As above, we randomly selected 100,000 sites along the reference genome spaced at least 500bp apart, and at these sites we created short insertions or short deletions in the reads. Where possible, the read edits were made in phase with a parental allele, otherwise numbers were chosen to match the empirical distribution. We then used platypus to detect indels in the simulated data and used the same filtering strategies as we did on the original data. However, when we compared the filtered results with the synthetic indels we created, we relaxed the match conditions such that indel length need not be correctly inferred, and indel position could fall within 5bp of the expected position. The number of callable sites was then estimated as for SNMs above.

Finally, to further verify our approach to SNM detection, we applied our filtering pipeline to the rhesus macaque trio data analysed by the recent ‘Mutationathon’ study (Bergeron, et al. 2022), and compared our results with those from five other research groups. As the candidate macaque mutations had been validated by PCR, they provide an opportunity to test the sensitivity of our pipeline. However, as only one F1 offspring was available in this test dataset, substantially reducing the power to distinguish false-positive mutations, we narrowed the acceptable allelic balance from (0.2 – 0.8) to (0.35 – 0.65).

### 2.5 Recombination detection

To infer crossover recombination in the parents, we used VCFtools to select biallelic SNPs with minimum genotype quality of 20 in each family and then applied ‘phase_by_transmission’ (window size = 100 heterozygous sites) from the Python package *scikit-allel* (version 1.3.5) to phase the two haplotypes in each offspring (Miles, et al. 2017). We visualised the haplotypes of informative SNPs in the offspring using R, and manually examined them to identify potential phasing errors in the parents. For example, the maternal haplotypes on 2R in fam27 of *D. simulans* displayed multiple identical breakpoints, consistent with mis-phasing of the parent (**supplementary fig. S6**), as did maternal haplotypes on the X chromosome in fam28 of *D. simulans*. We resolved a small number of such errors manually, by switching the parental phase at the shared breakpoint.

The resulting phased genomes should permit a direct count of crossover breakpoints, and thus an estimate of the recombination rate. However, substantial ‘noise’ was visible in the phasing at a scale of hundreds to thousands of bases (**supplementary fig. S6**). This noise could reflect sequencing errors, genotyping errors, or short phasing errors, but may also reflect gene-conversion events (Comeron, et al. 2012), which are difficult to robustly infer using short-read data. In principle, they might also represent very close pairs of crossover events, but because crossover interference is likely to suppress very close crossovers (Miller, et al. 2016), these can be excluded by setting a minimum threshold distance. To choose this threshold, we made use of the fact that recombination is generally absent in male *Drosophila* (Morgan 1912). As expected, we saw no compelling evidence of crossover recombination in males, but we did see a small number of short (i.e. ‘noisy’) recombinant haplotypes that could represent phasing errors or gene-conversion tracts. The longest tract was a region on 2L in a *D. simulans* individual that was supported by 1190 SNP markers. We therefore chose this as a threshold for the detection of true crossovers, on the basis that it was the longest (presumably erroneous) recombinant region detected in non-recombining males. When applied to the maternal haplotypes, this threshold allowed us to exclude 31,123 likely erroneous short haplotype blocks (median length 2 Kbp, 95% quantile = 188Kb).

Finally, given the potential for suppression of recombination by genomic inversions in heterozygotes, we used inversion-specific markers in *D. melanogaster* to detect inversions in the parental samples and examined whether the inversions, if present, affect the occurrence of recombination (Kapun, et al. 2014). The seven canonical inversions we examined were In(2L)t, In(2R)Ns, In(3L)P, In(3R)C, In(3R)K, In(3R)Mo, and In(3R)Payne.

### 2.6 Detection of transposable element insertions

The robust detection of TE insertions into repetitive regions is extremely challenging, and rarely possible using short-read data alone. However, a number of approaches can reliably detect TE insertions into low-complexity and gene-rich regions (Ewing 2015). We used TEFLoN (Adrion, et al. 2017) to identify TE insertions in our data, and we detected new insertions by comparing the TE locations between parents and offspring. TEFLoN first creates a pseudo-reference genome with TE sequences removed, then maps paired reads to the pseudo-reference genome and known TE sequences, identifying breakpoints to classify the reads into three categories: ‘presence’ reads that are a soft clipped or have a mate aligning to a TE, ‘absence’ reads whose alignment spans the breakpoints, and ‘ambiguous’ reads that are uninformative. We provided TEFLoN with a TE database that integrated two recently published TE databases in *Drosophila*: ‘Manually Curated Transposable Elements’ (MCTE) for *D. melanogaster* (Rech 2021) and a *D. simulans* species complex database (Chakraborty, et al. 2021). The MCTE database was originally created by running REPET (v.2.5) TE denovo pipeline on 13 genomes of natural *D. melanogaster*, followed by manual curation (Rech 2021). The *D. simulans* database was originally created by REPET and merged with *Drosophila* Repbase, and has been applied to detect TEs in *D. simulans, D. sechellia*, and *D. mauritiana* (Chakraborty, et al. 2021).

To confidently identify novel TE insertions, we then followed a similar logic to our SNM and indel filtering above, using the following steps. First, all the samples in each family were required to have at least 10 informative reads at a putative insertion site (presence reads or absence reads). Second, there could be no ‘presence’ reads in either parent. Third, the allelic balance of ‘presence’ reads in the focal offspring had to be between 0.2 – 0.8 (i.e. heterozygous). Fourth, the number of ‘presence’ reads in non-focal offspring could be no greater than 1, and no more than two individuals could have any reads supporting the ‘presence’ of the insertion. Finally, the candidate insertion could not be present in other families. For the candidate insertions on the X and Y chromosome in males, we halved the required number of informative reads, and required that the frequency of presence reads was greater than 0.8. We manually verified all the candidate insertions using IGV as we did to the SNMs.

To estimate the transposition rate per parental element, we counted the number of TEs in both parents of each family, requiring that each parental copy should have at least 10 informative reads, including at least 2 presence reads. The candidate TE was considered heterozygous if its frequency was in the range 0.2 – 0.8, and homozygous if the frequency was greater than 0.8. TEs on the X and Y chromosomes in the father were counted as one copy if the frequency was greater than 0.8.

### 2.7 Statistical analyses

We used binomial generalised linear mixed models to test for rate differences between populations and sexes and to quantify potential rate variation among individuals, using the Bayesian mixed-model R-package MCMCglmm (version 2.32; Hadfield 2010). To do this, we treated mutations (or recombination breakpoints, or insertions) as outcomes from a Bernoulli trial, with the estimated number of callable sites (or equivalent) as the number of trials. This approach naturally accounts for variation in power across the experiment (genome size, coverage, family size) and provides a robust and well-established framework for statistical testing. By transforming estimates of the latent rate parameter back on to the data scale, it also provides direct estimates of the rates and their credibility intervals.

To quantify differences in mutation rate among our three ‘populations’ (i.e. West African *D. melanogaster*, European *D. melanogaster, D. simulans*; termed ‘POPULATION’ in the model) and between the two parental sexes (‘SEX’), we fitted these terms as fixed effects. To quantify the variation among parents, we fitted parental id (‘PID’) as a random effect. MCMCglmm also fits a residual variance, such as might be caused by differences among observations of offspring that are not attributable to parents. A small number of mutations (or insertions) could not be assigned to their parent of origin, and we arbitrarily assigned half of these as maternal and half as paternal. This assignment will downwardly bias the difference between the sexes and the variance among parents, and in alternative models that are presented in supplementary material we separately explored the impact of (i) assigning the un-phased mutations differently, (ii) fitting chromosome arm as a fixed effect, and (iii) combining the parental and residual variance as a single term (**supplementary fig. S5**). For each model, we set the number of MCMC iterations to 50100000 with a thinning interval of 10000 and a burn-in of 100000 steps to allow the chain to reach stationarity. The MCMCglmm syntax for our full preferred model was:

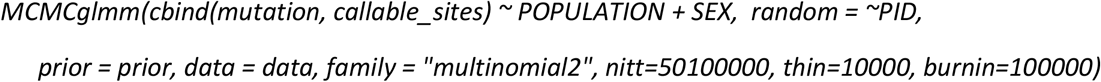

To improve mixing, we used the following diffuse parameter-expanded prior (Hadfield 2014):

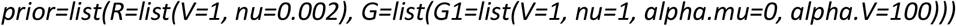

We similarly used MCMCglmm to estimate the recombination rate and to quantify variation in the recombination rate, with the following differences. First, data from males was excluded and parental sex was not fitted, because recombination is absent in male *D. melanogaster* and *D. simulans*. Second, chromosomal arm identity was included as a fixed effect, as recombination rates are known to vary between chromosome arms in *Drosophila* (Comeron, et al. 2012). Third, the number of Bernoulli trials was taken to be the genome length rather the number of ‘callable sites’, as recombination events would be detectable at any base within the spanned range, not just those at which sufficient coverage was available.

For transposable elements, we considered transposition rate in two contexts, transposition rate per base and transposition rate per parental element; the first used the length of host genome size that past depth filtering as a proxy for ‘callable sites’, and the second used the inferred copy number of TEs in host genomes to characterise the transposition rate.

## 3. Results

We sequenced parents and offspring of six unrelated outbred sibships from a West African population of *D. melanogaster*, a European population of *D. melanogaster*, and a European population of *D. simulans*. Each family comprised either a wild-collected or first-generation father together with an unrelated first-generation mother and five of their male F1 offspring. In total, we generated 9.5 billion 150bp paired-end Illumina sequencing reads, and after removing duplicates, the median genome coverage was above 30-fold for all but one of the 125 flies (range 28-108 fold; **supplementary fig. S1**). Based on the simulation of synthetic mutations in raw reads, we estimate the median proportion of callable sites was 83.1% (range 78.3-86.0%) for SNMs, and 78.3% (range 73.5%-80.1%) for short indels. Raw sequencing reads are provided in the European Genome-Phenome Archive with accession ID of PRJEB51956, and phased parental genomes are provided at https://doi.org/10.6084/m9.figshare.19733860.v1

Overall synonymous diversity (*π_s_*) was 1.27% in the West African *D. melanogaster*, 0.89% in the European *D. melanogaster* and 2.25% in the European *D. simulans*, similar to previously reported estimates (Parsch, et al. 2010; Campos, et al. 2017). The examination of a genomic relatedness matrix suggests that all the parents were unrelated except one pair from West African *D. melanogaster* (dmeA_15_F0 *vs*. dmeA_23_F0) and one pair from European *D. melanogaster* (dmeE_27_M0 *vs*. dmeE_30_F0), which were potentially third-degree relatives (i.e. first cousins; **supplementary fig. S2**). However, as these flies appeared in different families in this study, no further inbreeding occurred. Using breakpoint markers, we tested for the presence of seven chromosomal inversions in the two populations of *D. melanogaster*, and discovered twenty In(2L)t, three In(2R)Ns, six In(3L)P, six In(3R)K, and seven In(3R)Payne inversions, but no In(3R)C inversions segregating among the 48 haplotypes (**supplementary table S2**).

To test the performance of our pipeline for SNM identification, we applied the whole pipeline to trio data from rhesus macaques, as recently used by the ‘Mutationathon’ study (Bergeron, et al. 2022). Of the 33 SNMs there validated by PCR, our pipeline recovered 27, which was more than the other presented pipelines (**supplementary fig. S3**). Of the six false-positive SNMs that were mistakenly identified by some other pipelines, we only identified one. However, our pipeline also identified a further six candidates that have not been examined by PCR. Thus, at least in terms of sensitivity, our pipeline exhibits a performance at least as good as equivalent state-of-the-art approaches.

### 3.1 Mutation rates differ between populations and sexes

In total, we identified 58 SNMs across an estimated 17.45 billion callable sites, giving a crude overall *de novo* mutation rate across sexes and populations of 3.32 × 10^−9^ site^−1^ gen^−1^, with 95% binomial bounds of 2.52 - 4.30 × 10^−9^ (**fig. 1, supplementary figure S4**, **supplementary table S1**). However, the number of mutations varied among populations, sexes, and individuals (analysis below). Overall, SNMs were significantly biased toward transitions (overall 60.3 %, binomial *p*-value = 3.31 × 10^−5^; **fig. 1B**), but the mutational spectrum did not vary among populations (Fishers exact test; *p*-value = 0.82). We also identified six short deletions and three insertions, giving an overall short-indel rate of 5.00 × 10^−10^ site^−1^ gen^−1^, with 95% binomial bounds of 2.29 – 9.50 × 10^−10^ (**table 1**). Unlike the SNMs, the number of indels did not appear to differ markedly between populations, although they did appear to be biased towards males, with six of the attributable indel mutations occurring in the father.

**Figure 1:**
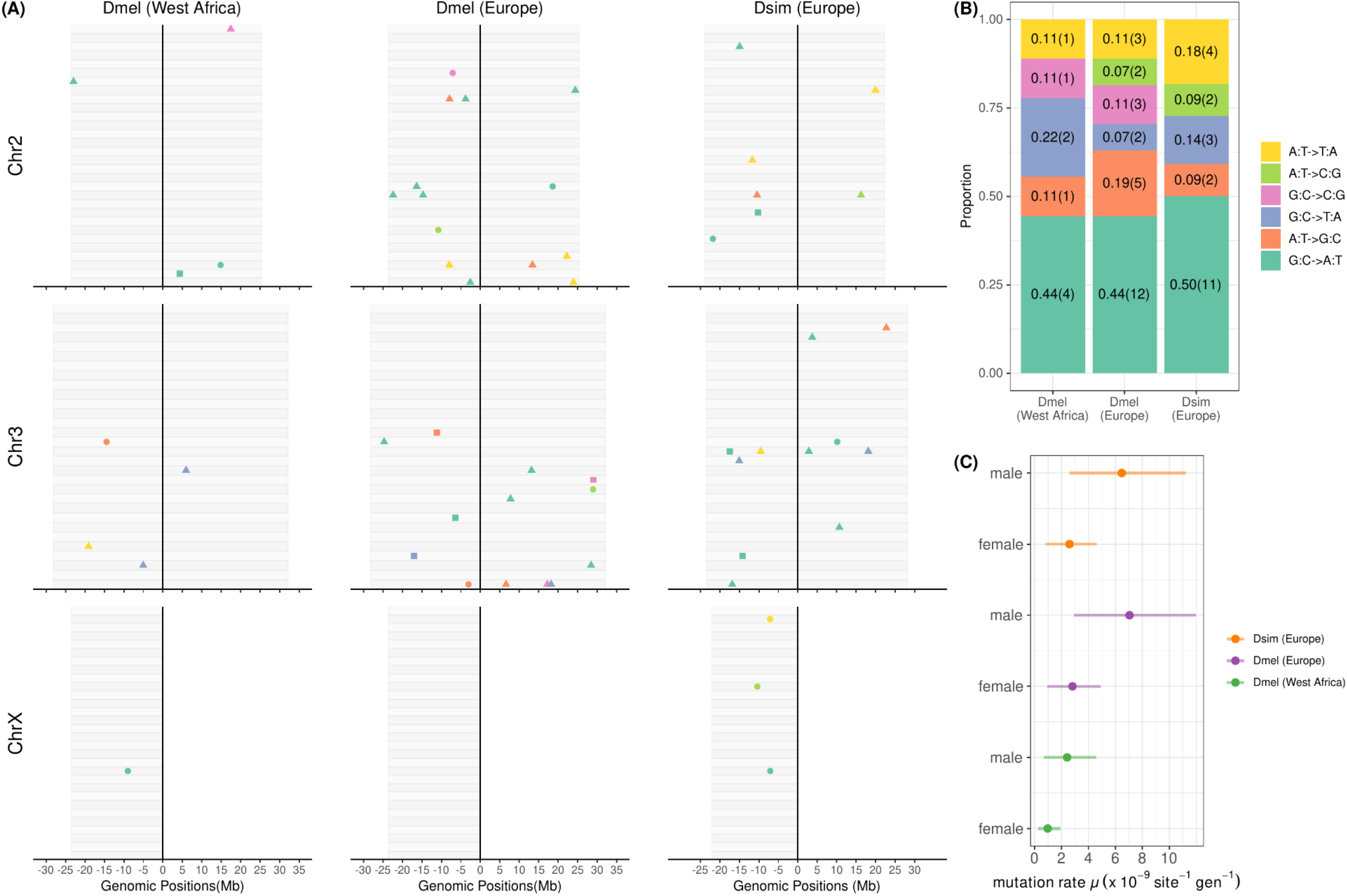
A summary of *de novo* SNMs identified in a West African and a European population of *Drosophila melanogaster*, and a European population of *Drosophila simulans*. (A) The genomic positions of the SNMs on chromosomes. Each grey bar represents the genome of one offspring. Point colour represents mutation type and point shape represents the parental origin of the mutation (triangle: paternal; circle: maternal). Square points are used to denote the SNMs with unknown parental origin due to the lack of informative surrounding SNP marker. (B) The SNM spectrum. The y-axis shows the proportion of each mutation type, and the numbers in brackets are the counts. (C) The mutation rate of SNMs and 95% CI estimated from our main Bayesian GLMM in which ‘population’ and ‘sex’ were fixed effects and parental ID was a random effect.

**Table 1.**
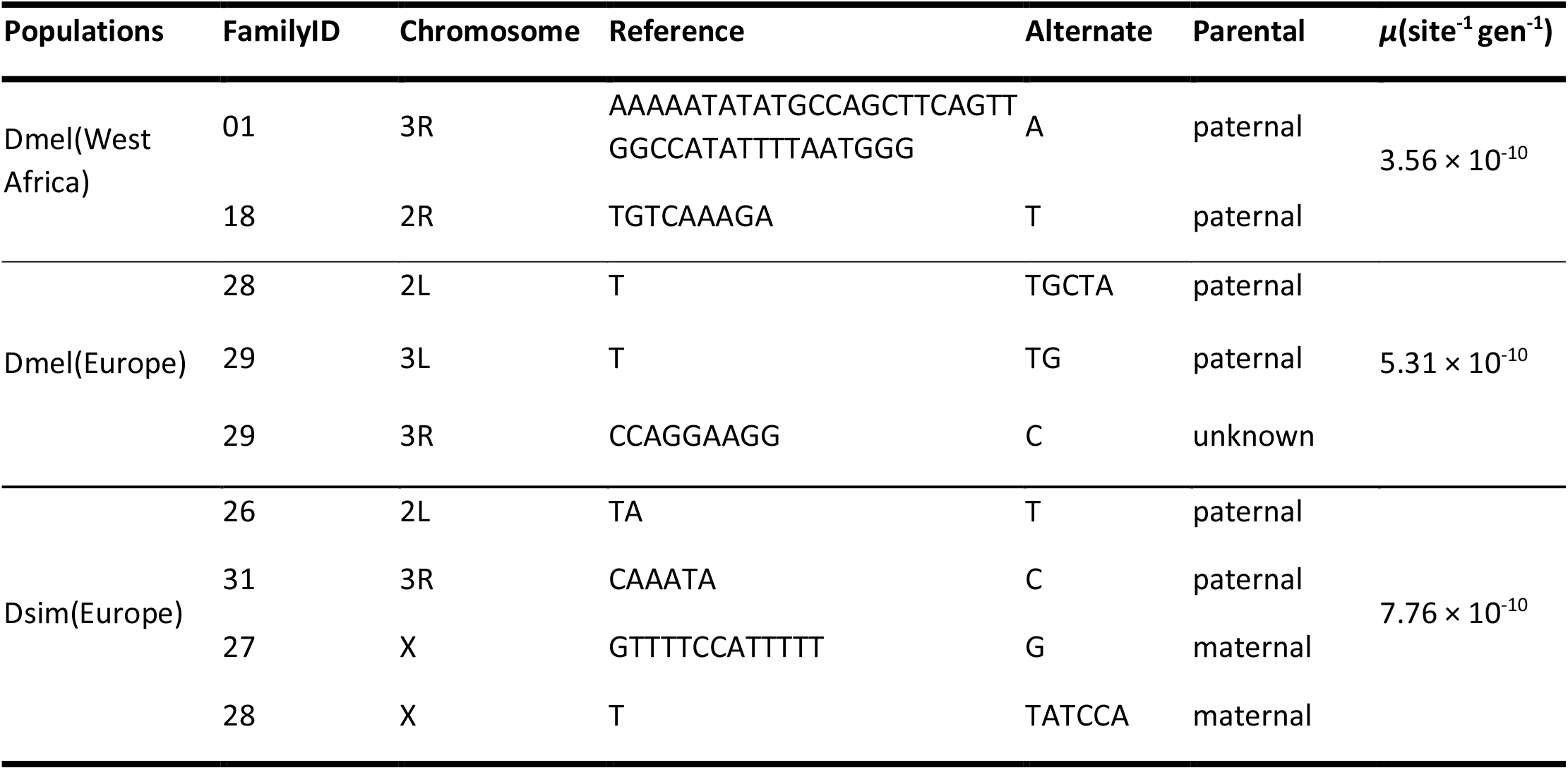
The summary of indel mutations.

Where the larger number of SNMs permitted, we used a Bayesian binomial linear mixed model to analyse variation in the mutation rate, fitting sex and population as fixed effects and parental identity as a random effect. Some mutations could not be attributed to one parent or the other, and we chose to assign half of these to males and half to females (underestimating the difference between sexes). Our model estimates are presented in **fig. 1C**, and alternative models are presented in supplementary material (**supplementary fig. S5**), although the results were not qualitatively different. Throughout, we treat the proportion of the posterior density overlapping zero a ‘p-value’, and we make no correction for multiple tests. We found the SNM rate of the West African *D. melanogaster* was significantly lower than that of the European populations of *D. melanogaster* or *D. simulans*, at 1.67 × 10^−9^ (95% HPD CI 0.54 - 3.14) *vs*. 4.86 × 10^−9^ (2.11 - 8.02) and 4.51 × 10^−9^ (1.94 - 7.75) respectively (*p*-value = 0.035, *p*-value =0.048; no correction for multiple testing), but that the European *D. melanogaster* and *D. simulans* populations did not differ significantly from each other (*p*-value = 0.863). Overall, we found that the mutation rate was higher in males than females, at 5.24 × 10^−9^ (2.96 - 7.83) *vs*. 2.05 × 10^−9^ (9.44 - 3.26) (*p*-value = 0.010)—although this was no longer significant if all of the un-phased mutations were assigned to females, thereby minimising the possible difference (*p*-value = 0.108). Despite an apparently striking range in the number of mutations seen per parent, after accounting for the fixed effects we found no strong support for substantial variation in mutation rate among individuals (65.48% of the remaining variance being attributable to among-individual variation, but with a 95% credibility interval of 5.00 - 99.96%).

### 3.2 Recombination rates vary among populations and chromosomes

We phased the parental haplotypes using the offspring data, and then used phase switches in the offspring to infer meiotic crossover breakpoints. Preliminary examination of the haplotype plots (**supplementary fig. S6**) identified a small number of large-scale phasing errors in the parents that were detectable through multiple sibling F1 individuals sharing identical breakpoints, for example on the X chromosome in the European *D. melanogaster* ‘fam30’, 2R in ‘fam27’, X in ‘fam28’ and 2R in ‘fam29’ in the European *D. simulans*. We manually corrected these errors by switching the parental phases, and the correctly phased parental genomes are available at figshare.com (https://doi.org/10.6084/m9.figshare.19733860.v1). Consistent with an absence of recombination in male *Drosophila* (Morgan 1912), no recombinant haplotypes strongly supportive of crossover were detected in males. However, we did identify shorter apparent recombinant regions with maximum length of up to ~1Mbp in males and ~4Mbp in females (corresponding to 1190 SNP markers), which we believe represented genotyping errors (longer regions) or gene conversion events (shorter regions). After excluding these regions, we identified 103 recombination events in female West African *D. melanogaster*, 74 in European *D. melanogaster*, and 109 in European *D. simulans* (**fig. 2**, **supplementary table S2**, **supplementary fig. S4**). This corresponds to 3.43, 2.55 and 3.63 detectable crossovers per individual, resulting in raw recombination rates of 2.59 cM/Mb (95% binomial confidence interval: 2.11 - 3.14), 1.92 cM/Mb (1.51 - 2.42), and 3.03 cM/Mb (2.49 - 3.66), respectively. To examine the robustness of our inference, we explored the impact of marker thresholds lower than 1190 and found that the number of recombination events remain stable even if we varied the threshold (**supplementary fig. S7**). We could not detect any correlation between the number of crossovers detected and the number of available SNP markers in a chromosome arm (*p*-value = 0.1264, **supplementary fig. S8**), suggesting our ability to detect crossovers is not strongly affected by variation in the number of SNPs.

**Figure 2:**
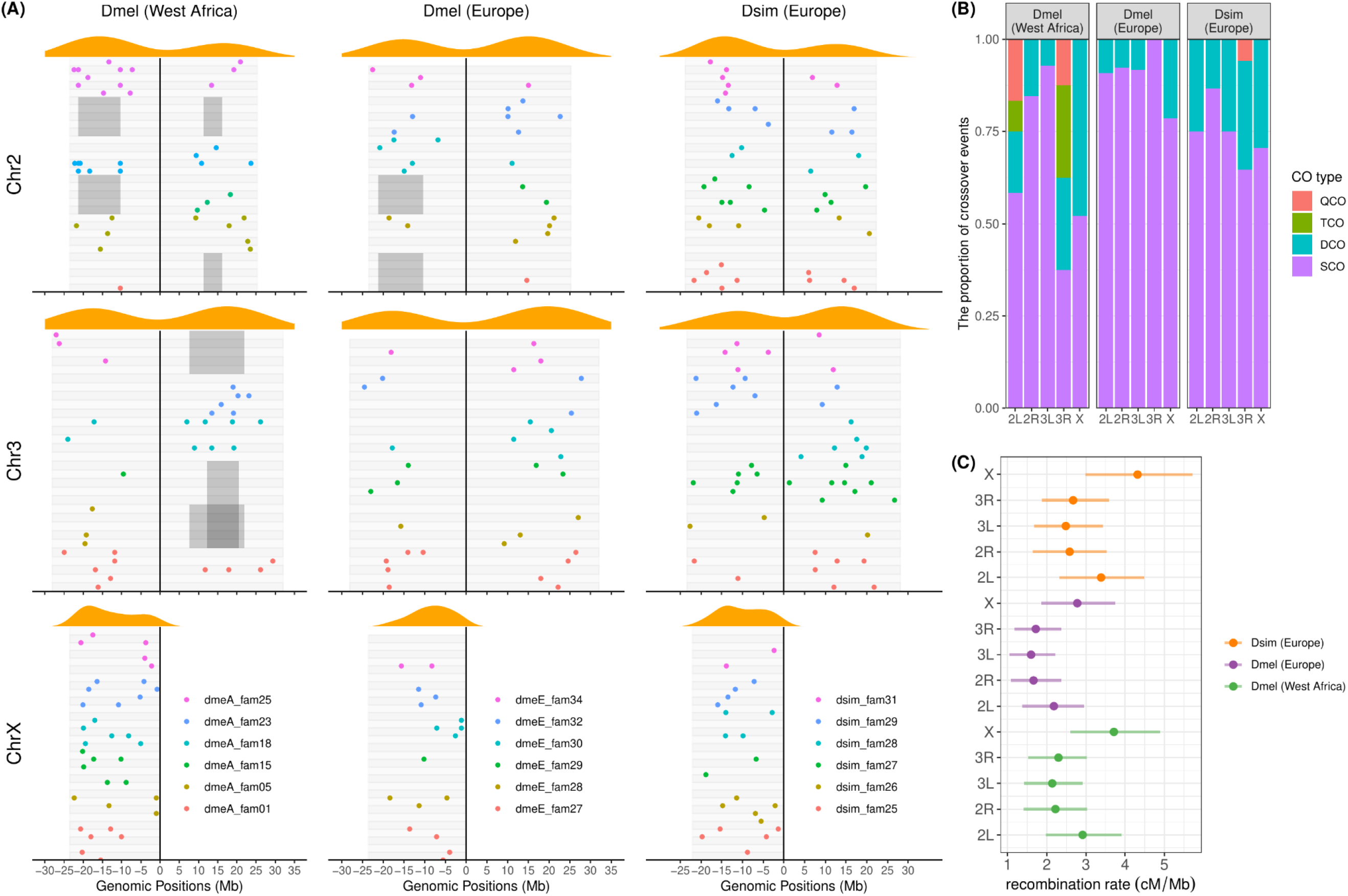
A summary of recombination (crossovers) in the females of three populations of *Drosophila*. (A) The genomic positions of breakpoints on chromosomes. Each grey bar represents an offspring individual and colours denote the breakpoints in different families. Orange density plots above represent the density of breakpoint along each chromosome. The classical inversions of *D. melanogaster* (In(2L)t, In(2R)NS, In(3R)K and In(3R)Payne) heterozygous in the parents are shown as grey areas. (B) The proportion of crossover types; SCO: one crossover event on a chromosomal arm during one meiotic process; DCO: two crossover events in one meiotic process; TCO: three crossover events; QCO: four crossover events. (C) The recombination rate and 95% CI estimated from a Bayesian GLMM in which ‘population’ and ‘chromosome’ were fixed effects and parental ID was a random effect.

As expected, there were few recombination breakpoints close to centromeres and telomeres. In *D. melanogaster*, the nearest breakpoint to a centromere was identified at a distance of 6.8Mb on 2L (**fig. 2A**). In *D. simulans*, all the breakpoints were >3.7Mb away from centromere except one identified at 1.3Mb to the centromere on 3R. Among the crossovers, we observed 45 double crossovers, where two crossovers were detectable on the same chromosomal arm in one meiosis, with 18 in West African *D. melanogaster*, 6 in European *D. melanogaster* and 21 in European *D. simulans*. Double crossovers were particularly common for the X chromosome in West African *D. melanogaster*, where nearly half of the crossovers occurred in doubles, and there were three triple crossovers and three quadruple crossovers (**fig. 2B**).

To quantify maternal recombination rate more robustly, and to test if recombination rates varied among populations, individuals, and chromosome arms, we applied MCMCglmm using models analogous to those used above for mutation rate. We fitted population and chromosome arm as fixed effects, and individual as a random effect. In addition, because large inversions are expected to suppress recombination, we fitted a fixed effect of ‘inverted’ for maternal chromosome arms in individuals inferred to be heterozygous for any of the seven large inversions we examined. In the absence of large inversions, we found that the recombination rate in flies from the West African population of *D. melanogaster* was significantly higher than that in flies from the European populations of *D. melanogaster* at 3.44 (95% HPD CI 2.72 – 4.18) cM/Mb *vs*. 2.06 (1.57 - 2.57) cM/Mb (*p*-value = 0.004), but not higher than the European *D. simulans* 3.04 (2.45 - 3.73) cM/Mb (*p*-value = 0.430). The European *D. melanogaster* had a lower recombination rate than *D. simulans* (*p*-value = 0.019). We did not see any recombination on chromosomal arms that were heterozygous for large classical inversions in *D. melanogaster*, and the presence of an inversion significantly suppressed recombination (*p*-value < 2 × 10^−4^). Using a threshold *p*-value of 0.05 without correcting for multiple testing, we detected a significantly higher recombination rate on arm 2L (3.62[2.71 - 4.63] cM/Mb) and the X chromosome (3.57[2.74 - 4.46] cM/Mb) than that on arms 2R (2.43[1.70 – 3.20] cM/Mb) and 3L (2.05[1.49 - 2.67] cM/Mb). The recombination rate on arm 3R was estimated at 2.70 [2.00 - 3.47] cM/Mb) and was not significantly different from other chromosomal arms. After accounting for all of the fixed effects, we found little evidence for variation among individuals (48.6% of the remaining variance being attributable to among individual variation; 95% CI of 0 – 96.2%).

### 3.3 TE insertion rates differ between populations and sexes

We used TEFLoN (Adrion, et al. 2017) to annotate TEs in parents and offspring, and we identified new insertions as those that were heterozygous in a single F1 offspring, but absent from all other individuals in the study. In total, TEFLoN identified *ca*. 200,000 TE copies across the 24 parental *D. melanogaster* genomes. The 15 most abundant TEs in the parents are shown in **supplementary fig. S9**. As expected, the rankings of TE abundance were very similar between the two populations of *D. melanogaster*, with *Gypsy, Helitron* and *Jockey* elements being the most abundant, followed by *Pao, P, TcMar-Tc1, CR1, R1, CMC-Transib*, and *INE-1*. In total we identified *ca*. 138,000 TE copies across the 12 *D. simulans* parents, i.e. about two thirds of that in *D. melanogaster*. The ranking was slightly different in the *D. simulans* population, as predicted from the known differences in the TE community between these species (Merel, et al. 2020), with *Helitron* elements being the most common TE, followed by *Gypsy* and *Jockey*. DNA transposons made up 39% of all TEs in *D. melanogaster* and 44% in *D. simulans*; LTRs made up 36% and 34% in the two species, respectively; non-LTRs made up 25% and 22%, respectively. In all populations, most individual TE insertions were rare, with 45.7% being singletons and 17.8% being doubletons, and in general retrotransposons were more frequent than DNA transposons.

By comparing the parents and offspring in each family, we identified 89 new TE insertions across the three population samples, with 18 in West African *D. melanogaster*, 45 in European *D. melanogaster*, and 26 in European *D. simulans* (**supplementary fig. S4**). Although these numbers probably represent lower bounds for the true number of insertions, as those falling in repetitive hard-to-map regions or within other TEs are unlikely to be detectable, our estimates are likely to be a good estimate of the rate of transposition into non-repetitive, gene-rich, euchromatin. The 18 new insertions in the West African *D. melanogaster* came from just 2 TE super-families: *CMC-Transib* and *Gypsy*, with the largest contributor being *CMC-Transib* (17 insertions) (**fig. 3A**, **supplementary fig. S10**). The active super-families appeared to be strikingly different in Europe (Fisher’s exact test *p*-value = 0.0005), covering 5 super-families with 19 *Gypsy*, 16 *CMC-Transib*, 6 *Pao*, 3 *Jockey* and 1 *hAT-hobo* insertions. The highly active *CMC-Transib* elements in *D. melanogaster* (3.09 × 10^−3^ copy^−1^ gen^−1^) were less active in *D. simulans*, only showing 3 insertions with an insertion rate of 5.67 × 10^−4^ copy^−1^ gen^−1^. We observed 6 *Copia* insertions and 8 *TcMar-Mariner* insertions in *D. simulans*, but none in *D. melanogaster*. The insertion rates for *Copia* and *TcMar-Mariner* in *D. simulans* were estimated as high as 2.71 × 10^−3^ copy^−1^ gen^−1^ and 3.06 × 10^−2^ copy^−1^ gen^−1^, respectively. We did not observe new *Jockey* insertions in *D. simulans*. However, underlying these raw counts is substantial variation among individuals, for example, all 11 *Nomad* insertions (*Gypsy* super-family) in European *D. melanogaster* occurred in the same individual parent. There were also apparent differences between the sexes, as among those TE insertions that could be attributed to the parent of origin in *D. melanogaster* the majority occurred in males (38 *vs*. 15), whereas in *D. simulans* males and females had a similar number of TE insertions (12 *vs*. 11).

**Figure 3:**
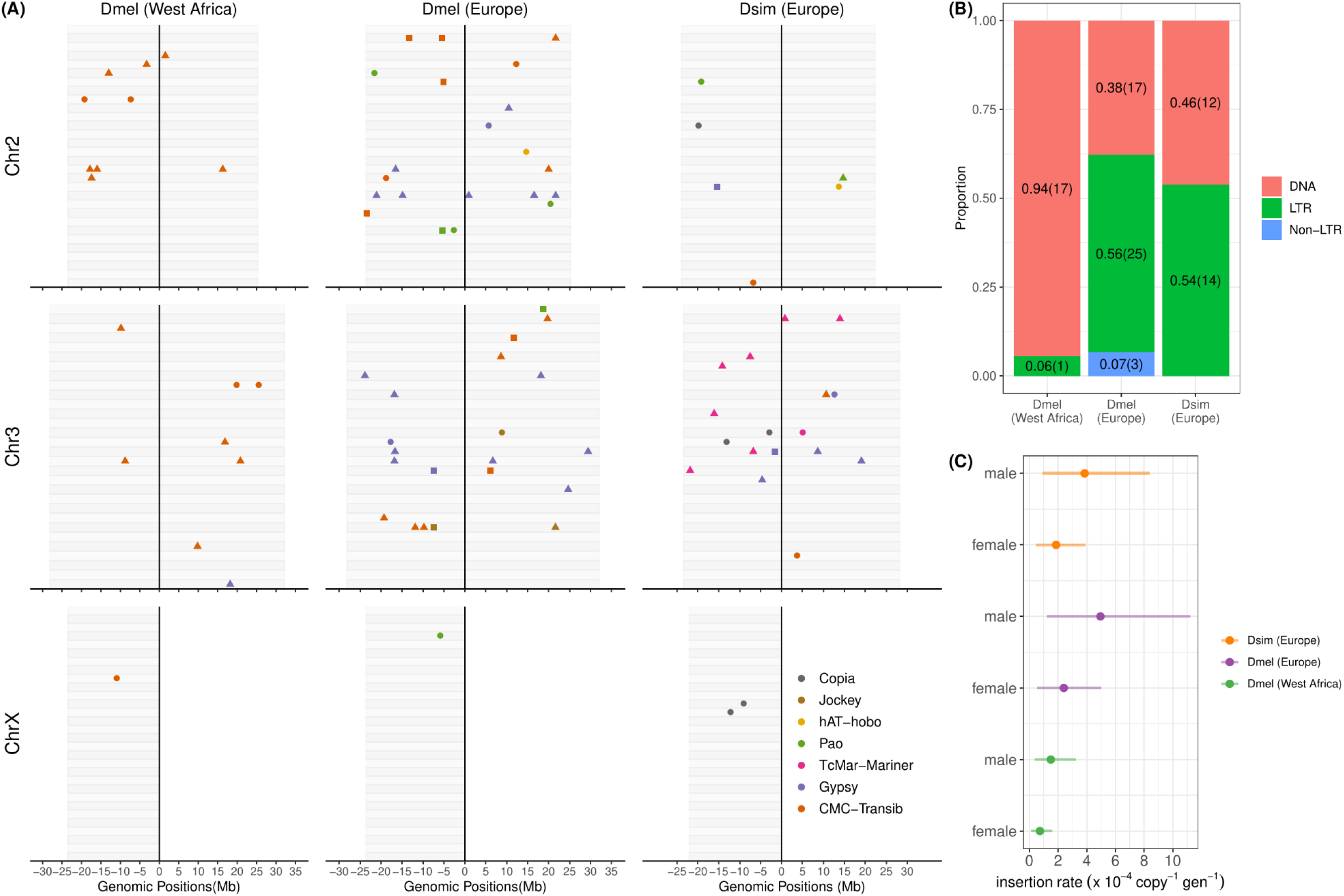
A summary of TE insertions in the three populations of *Drosophila*. (A) The genomic positions of the new TE insertions on chromosomes. Each grey bar represents an offspring individual, point colour represents TE superfamilies, and point shape represents the parental origin of the insertion (triangle: paternal; circle: maternal; square: unknown). As above, insertions with unknown parental origin are shown as squares. (B) The proportion of new insertions of DNA elements, LTRs and non-LTRs in the three populations. (C) The insertion rate (per copy per generation) and 95%CI estimated from a Bayesian GLMM where ‘population’ and ‘sex’ were fixed effects and parental ID was a random effect.

As above, we used MCMCglmm to quantify transposition rates and their variation between populations and sexes. Transposition rates can be quantified relative to the number of TE copies in the parent or relative to the number of sites available in the genome, depending on whether one is interested directly in the insertion activity of TEs or instead in their impact on the genome. Accordingly, we fitted models with either the number of parental element copies as the ‘trials’ in the binomial model, or the number of callable sites. Similar to the SNM rate, insertion rates were significantly lower in West African *D. melanogaster* than in European *D. melanogaster*, both when quantified per parental element (*p*-value = 0.026) (**fig. 3C**) and per base (*p*-value = 0.035). The European population of *D. simulans* had an intermediate rate that was not significantly different from the two *D. melanogaster* populations. Males tended to have slightly more insertions than females, but this effect was not significant (*p*-value = 0.118 for copy-based rate, *p*-value = 0.071 for base-based rate). Overall, per parental copy, we estimate 8.99 × 10^−5^ (5.33 - 14.21) insertions copy^−1^ gen^−1^ for the West African population *vs*. 23.36 × 10^−5^ (17.04 - 31.26) insertions copy^−1^ gen^−1^ in the European *melanogaster*, and 18.70 × 10^−5^ (12.25 - 27.48) insertions copy^−1^ gen^−1^ in the European *simulans* (**supplementary table S3**). Per callable base, the rates were 2.95 × 10^−9^ (1.73 - 4.62) site^−1^ gen^−1^, 7.54 × 10^−9^ (5.50 - 10.09) site^−1^ gen^−1^ and 4.38 × 10^−9^ (2.86 - 6.42) site^−1^ gen^−1^ for the three populations, respectively.

## 4. Discussion

Here we used parent-offspring sequencing to directly estimate the mutation, recombination, and transposition rates in two populations of *D. melanogaster* and one population of *D. simulans*. By crossing unrelated wild-collected or first generation flies we were able to avoid many of the biases that can arise from inbreeding in mutation-accumulation and mapping studies, such as selection against recessive deleterious mutations or the direct impact of new mutations on mutation, recombination, or transposition rates. In addition, because we could identify the parent of origin for the majority of events, we were able to quantify differences in rate between the sexes and between individuals.

### 4.1 Overall mean rates of mutation, recombination, and transposition

Previous estimates of the *de novo* SNM rate for *D. melanogaster* span the range from 2.8 to 6.0 × 10^−9^ site^−1^ gen^−1^ (Keightley, et al. 2014; Sharp and Agrawal 2016), and our overall mean estimate of 3.3 × 10^−9^ site^−1^ gen^−1^ across *D. melanogaster* and *D. simulans* is in close agreement. This is at the lower end of the range seen for metazoa, which range from 2.0 to 16.6 × 10^−9^ (Yoder and Tiley 2021), but is very similar to other insects, including bumblebees at 3.6 × 10^−9^ (Liu, et al. 2017), honeybees at 3.4 × 10^−9^ (Yang, et al. 2015), and butterflies at 2.9 × 10^−9^ (Keightley, et al. 2015). Our estimates correspond to an average of just 0.8 new SNMs present in each new female embryo.

As with previous studies, we observed a mutational spectrum that is biased toward transitions, particularly G:C->A:T (Schrider, et al. 2013), and we observed one complex mutational event, with two mutations occurring only 2bp apart. This one event out of 57 is consistent with a previous MA study of two *D. melanogaster* lines that suggested around 2% of SNMs in *Drosophila* are complex (Schrider, et al. 2013), likely as result of the error-prone polymerases utilized to bypass some DNA lesions during synthesis (Segurel, et al. 2014). Compared to SNMs, new short indels are scarce in *Drosophila*, and we only detected 9 in total. This implies a rate of 0.5 × 10^−9^ site^−1^ gen^−1^, which is about half that estimated by Keightley et al (2014) who estimated 1.2 × 10^−9^, but similar to the estimate of Schrider et al. (2013) at *ca*. 0.4 × 10^−9^. Indel rates vary dramatically across multicellular eukaryotes, from 0.3 × 10^−9^ site^−1^ gen^−1^ in mouse to 1.8 × 10^−9^ site^−1^ gen^−1^ in humans (reviewed in Sung, et al. 2016), placing *Drosophila* at the lower end of the range.

Recombination rates are known to vary dramatically among taxa, from 0.03 cM/Mb to 28.10 cM/Mb among multicellular plants and animals (Stapley, et al. 2017). Although our parent-offspring approach lacks power, our raw estimate of the overall recombination rate in females is 2.50 cM/Mb, very close to that from a previous study of *D. melanogaster* of *ca*. 2.45 cM/Mb (Comeron, et al. 2012). As expected, recombination events appeared to be suppressed near to centromeres, and the presence of large chromosomal inversions significantly reduced the recovery of recombinants. Previous studies have reported the variation of recombination rate between chromosomes, for example the higher rate on X chromosome in African *D. melanogaster* (Chan, et al. 2012), which is in line with our study after excluding the impact of large chromosomal inversions.

Although TE insertions provide an unusually disruptive form of mutation (Wells and Feschotte 2020), transposition rates are more difficult to estimate directly and have not been widely studied. Here we estimate an overall rate of at least 16.7 × 10^−5^ insertions per parental TE copy per generation corresponding to 4.93 × 10^−9^ insertions per site per generation, or approximately one new insertion in each new female embryo. Our estimates are 2.3-fold higher than a previous estimate (Adrion, et al. 2017) in *Drosophila* after ~150 generations mutation accumulation (2.11 × 10^−9^ site^−1^ gen^−1^). Our estimate of the TE insertion rate is *ca*. 1.5-fold higher than the rate of *de novo* SNMs. This suggests that the impact of deleterious mutation on the genome from TE insertions is likely to be substantially higher, especially given their likely effects, than the impact of deleterious SNMs.

### 4.2 Rate differences between populations, sexes, and individuals

Underlying these rates of mutation, recombination, and TE insertion was substantial variation among populations, sexes, and individuals. We observed an SNM rate that was nearly 3-fold (95% HPD CI: 1.07 – 6.58) higher in the European population than in the West African population of *D. melanogaster* (4.8 *vs*. 1.6 × 10^−9^ site^−1^ gen^−1^), with European *D. simulans* being more similar to European *D. melanogaster* (4.5 *vs*. 4.8 × 10^−9^ site^−1^ gen^−1^). It is interesting to note that our low estimate for West African *D. melanogaster* is close to that of the only other African estimate (2.8 × 10^−9^ site^−1^ gen^−1^)— which was similarly based on parent-offspring sequencing from a single family collected in Ghana (Keightley, et al. 2014)—whereas our estimate for European *D. melanogaster* is closer to those from inbred North American isolates (Schrider, et al. 2013; Huang, et al. 2016; Assaf, et al. 2017).

Remarkably, we also saw a substantial (2.5-4.3 fold) difference in the SNM rates between male and female flies, approaching that seen in primates (3-8 fold; Taylor, et al. 2006; Arnheim and Calabrese 2009; Segurel, et al. 2014). Direct observation of sex differences in mutation rate has not been reported previously for *Drosophila*, as other studies have not assigned mutations to the parent of origin. However, Schrider et al (2013) examined differences in mutation accumulation on the X and autosomes within MA lines, suggesting a potential difference between the two sexes, and a study of neo-sex chromosomes in *D. miranda* suggested twice as many mutations in males as females (Bachtrog 2008). Finally, although we did observe many more mutations in some individuals than in others (up to nine among the offspring of one *D. melanogaster* male, where the expected number was 3.9), and previous studies have suggested substantial differences between inbred lines (Schrider, et al. 2013), after accounting for the fixed effects of population and sex we had little power to quantify the level of among-individual variation (95% CI: 0 to 0.79). This is in contrast to an MA study on the green alga *Chlamydomonas reinhardtii* that found a sevenfold variation among the strains (Ness, et al. 2015), and the presence of hypermutation in some individual humans (Kaplanis, et al. 2021).

As expected, we saw no compelling evidence for crossovers in male *Drosophila* (Morgan 1912), but we did see some evidence for variation in female recombination rate between populations, with the model estimates for female African and European *D. melanogaster*, and European *D. simulans* of 3.4 cM/Mb, 2.1 cM/Mb and 3.0 cM/Mb, respectively. This is comparable to previous estimates from *D. melanogaster* (Comeron, et al. 2012; Miller, et al. 2016), and is consistent with Chen et al (2012) who suggested the North American *D. melanogaster* had a lower recombination rate than the African populations. Our estimate for *D. simulans* is in close agreement with the observation that the linkage map length in *D. simulans* is around 1.3-fold longer than that in *D. melanogaster* (True, et al. 1996).

As with mutation and recombination rate, we identified substantial variation among populations in the rate of transposition. Notably, the West African population of *D. melanogaster* had a 2.6-fold lower insertion rate than the European population, with the rate in the *D. simulans* population more similar to that of European *D. melanogaster*. We observed some TEs were highly active in specific samples, consistent with observations from the MA lines of *D. melanogaster* (Adrion, et al. 2017).

### 4.3 Causes and consequences of rate variation

Per-generation mutation rates are generally thought to be determined by DNA replication fidelity, DNA repair efficiencies, mutagen exposure, and the number of germline cell divisions per generation (Baer, et al. 2007). Consequently, sex and age are often found to be critical factors, and male mutation rates are usually several-fold higher than female rates in older primates (Taylor, et al. 2006; Arnheim and Calabrese 2009; Kong, et al. 2012; Segurel, et al. 2014; Thomas, et al. 2018; Kaplanis, et al. 2021). The difference between the sexes has historically been attributed to the larger number of cell divisions required in the production of sperm, but recently studies on humans suggests that DNA damage and maternal age were also critical in determining the germline mutations (Gao, et al. 2019). However, environmental factors such as temperature and exposure to mutagenic chemicals or ultraviolet radiation are also likely to play a key role—perhaps especially so in small terrestrial invertebrates (Pfeifer, et al. 2005; Baer, et al. 2007; Kaplanis, et al. 2021; Waldvogel and Pfenninger 2021). The differences we see here between males and females, and perhaps also between West African and European populations, might therefore be attributable to a higher male mutation rate *per se*, or to differences between wild-collected (European males) and first-generation laboratory flies (females and west African flies) in age, stress, or mutagen (UV) exposure.

Recombination rates in *Drosophila* also not only depend on genetic background, with up to a two-fold difference between North American genotypes (Hunter, et al. 2016), but also depend on female age, number of matings, nutrition, temperature, and pathogen exposure (Redfield 1966; Hunter and Singh 2014; Stapley, et al. 2017; Aggarwal, et al. 2021). However, as all of the female flies used in this experiment were first-generation laboratory flies and were thus similar in age and environmental exposure, the difference in recombination rates more likely resulted from genetic differences.

Like recombination, high rates of transposition rate have been associated with variation in temperature, irradiation, chemical agents, and pathogens (Capy, et al. 2000; Guerreiro 2012). The worldwide colonization of *D. melanogaster* and *D. simulans* exposed the immigrants to new environmental conditions—which has been claimed to reactivate TEs—and may have exposed them to invasion by new TEs from sympatric species (Guerreiro 2012; Schwarz, et al. 2021). All of these factors could come into play in explaining the variation we see here. However, most of the male flies used in this study were F0 wild-collected flies and some were F1, and maintained in the laboratory conditions at a constant temperature. Given the difference of UV radiation and temperature between laboratory and wild conditions, our results may not directly and accurately reflect the natural populations, but should be closer than many other studies that use laboratory inbred lines or MA lines (Roles and Conner 2008).

Regardless of the causes of this variation in mutation, recombination, and transposition, it has potentially important implications for studies of *Drosophila* evolution. Most notably, *D. melanogaster* mutation-rate estimates are widely used to derive estimates of *Ne* from measures of diversity (Kimura 1991), and to apply timescales to the evolutionary process in *Drosophila* (Obbard, et al. 2012; Kapopoulou, et al. 2018) and even beyond (Miles, et al. 2017). On the one hand, our finding of a broadly similar rate in European *D. melanogaster* and *D. simulans* is reassuring; it may suggest that rates are usefully transferable between species. On the other hand, the possibility of a nearly 3-fold difference in mutation rate between African populations and the out-of-Africa diaspora could have substantial implications for estimates of *Ne* or timescales of *Drosophila* history.

## Abbreviations

SNMs: single nucleotide mutations
SNVs: single nucleotide variants
TEs: transposable elements
indels: insertions and deletion mutations
cM: centiMorgan(s)
Mb: Mega base pair(s)

## Acknowledgments

We thank Mallam Adamu Shuaibu, and Keith and Sue Obbard for assistance in collecting the *Drosophila*, and thank Brian Charlesworth for encouraging us to examine the impact of inversions on recombination rate.

## Supplementary Figure Legends

**supplementaryFigureS1**: Boxplots of sequencing coverage of all samples. Coverage was calculated as a mean value for each 1000bp window along the genome. Outliers are not shown in the figure. ‘dmeA’: the West African *D. melanogaster*; ‘dmeE’: the European *D. melanogaster*; ‘dsim’: the European *D. simulans*.

**supplementaryFigureS2**: The relatedness of the flies collected from the wild. For each population, we made 6 families by mating a female (‘F0’) and male (‘M0’). Two pairs in the *D. melanogaster* population (dmeA_15_F0 *vs*. dmeA_23_F0; dmeE_27_M0 *vs*. dmeE_30_F0) may be potentially 3rd degree relatives (e.g. first cousins).

**supplementaryFigureS3**: The test of our SNMs pipeline on a trio dataset of rhesus macaques (Bergeron, et al. 2022), which includes 33 PCR-confirmed *de novo* mutations of which we recovered 27 (here in red). The other research groups are - LB: Lucie Bergeron; SB: Søren Besenbacher; TT: Tychele Turner; RW: Richard Wang CV: Cyril Versoza.

**supplementaryFigureS4**: The number of new variants in three *Drosophila* populations: the number of *de novo* SNMs (top panel), the number of recombination breakpoints (centre panel) and the number of TE insertions (bottom panel). For the SNMs and TEs, we used ‘X’ to indicate a new variant on the X chromosome and ‘Y’ on Y chromosome. Blue bars denote a variant originating in males; orange bars denote a variant originating in females; grey bars mean that the variation could not be assigned to one parent or the other due to lack of informative SNP markers.

**supplementaryFigureS5**: The mutation rate and 95% CI estimated using different datasets and different models. Depending on how we assign the un-attributable mutations we created three datasets. ‘D1’: half of these mutations to the males and half to females; ‘D2’: all these mutations to males; ‘D3’: all these mutations to females. ‘modelA’ includes ‘POPULATION’ and ‘SEX as fixed effects and parental ID ‘PID’ as random effect; ‘modelB’ incorporates ‘CHROM’ as an additional fixed effect based on modelA; ‘modelC’ excludes random effect ‘PID’ from the modelA.

**supplementaryFigureS6**: The haplotypes phased using informative SNP markers. The x-axis represents the SNP markers on chromosome arms. The y-axis represents the offspring in each family. Two colours are used to distinguish two different haplotypes in the mother or father.

**supplementaryFigureS7**: The relationship between the threshold of SNP markers to infer a haplotype block and its resulting number of breakpoints. The x-axis is the threshold of SNP markers which we used to infer a haplotype block. Any haplotype block containing SNP markers less than the threshold will be considered as ‘noisy’. The y-axis represents the total number of inferred breakpoints across 5 families in each population given the threshold. We didn’t manually check and correct a few potentially phasing errors in this figure.

**supplementaryFigureS8:** The relationship between the number of SNP markers aggregated by family and the number of crossovers on different chromosomes.

**supplementaryFigureS9**: The total number of TE copies by TE super-family in 12 parental samples for each population. Only the top 15 most abundant TEs are shown.

**supplementaryFigureS10:** The number of new TE insertions identified in each parental sample. We identified a total of 89 insertions and 13 of them were not attributable to the parents due to lack of informative SNP markers and not shown in the figure. ‘dmeA’: the West African *D. melanogaster*; ‘dmeE’: the European *D. melanogaster*; ‘dsim’: the European *D. simulans*. Sample names with ‘F0’ denote female (mother); ‘M0’ for male (father).

**supplementaryTableS1**: The number of SNMs and indel mutations in each family and their corresponding callable sites and estimated mutation rates.

**supplementaryTableS2**: The details of genomic inversions and recombination breakpoints.

**supplementaryTableS3**: The number of TE insertions and the corresponding insertion rate in each *Drosophila* family and TE superfamily.

## References

Adrion JR, Song MJ, Schrider DR, Hahn MW, Schaack S. 2017. Genome-Wide Estimates of Transposable Element Insertion and Deletion Rates in Drosophila Melanogaster. Genome Biol Evol 9:1329–1340.

Aggarwal DD, Rybnikov S, Sapielkin S, Rashkovetsky E, Frenkel Z, Singh M, Michalak P, Korol AB. 2021. Seasonal changes in recombination characteristics in a natural population of Drosophila melanogaster. Heredity.

Arnheim N, Calabrese P. 2009. Understanding what determines the frequency and pattern of human germline mutations. Nature Reviews Genetics 10:478–488.

Assaf ZJ, Tilk S, Park J, Siegal ML, Petrov DA. 2017. Deep sequencing of natural and experimental populations of Drosophila melanogaster reveals biases in the spectrum of new mutations. Genome Research 27:1988–2000.

Bachtrog D. 2008. Evidence for male-driven evolution in Drosophila. Molecular Biology and Evolution 25:617–619.

Baer CF, Miyamoto MM, Denver DR. 2007. Mutation rate variation in multicellular eukaryotes: causes and consequences. Nature Reviews Genetics 8:619–631.

Bergeron LA, Besenbacher S, Turner T, Versoza CJ, Wang RJ, Price AL, Armstrong E, Riera M, Carlson J, Chen HY, et al. 2022. The Mutationathon highlights the importance of reaching standardization in estimates of pedigree-based germline mutation rates. Elife 11.

Campbell CD, Chong JX, Malig M, Ko A, Dumont BL, Han LD, Vives L, O’Roak BJ, Sudmant PH, Shendure J, et al. 2012. Estimating the human mutation rate using autozygosity in a founder population. Nature Genetics 44:1277–1281.

Campos JL, Zhao L, Charlesworth B. 2017. Estimating the parameters of background selection and selective sweeps in Drosophila in the presence of gene conversion. Proc Natl Acad Sci U S A 114:E4762–E4771.

Capy P, Gasperi G, Biemont C, Bazin C. 2000. Stress and transposable elements: co-evolution or useful parasites? Heredity 85:101–106.

Chakraborty M, Chang CH, Khost DE, Vedanayagam J, Adrion JR, Liao Y, Montooth KL, Meiklejohn CD, Larracuente AM, Emerson JJ. 2021. Evolution of genome structure in the Drosophila simulans species complex. Genome Research 31:380–396.

Chan AH, Jenkins PA, Song YS. 2012. Genome-wide fine-scale recombination rate variation in Drosophila melanogaster. Plos Genetics 8:e1003090.

Charif D, Lobry JR. 2007. SeqinR 1.0-2: a contributed package to the R project for statistical computing devoted to biological sequences retrieval and analysis. In. Structural approaches to sequence evolution: Springer. p. 207–232.

Chuong EB, Elde NC, Feschotte C. 2017. Regulatory activities of transposable elements: from conflicts to benefits. Nature Reviews Genetics 18:71–86.

Comeron JM, Ratnappan R, Bailin S. 2012. The Many Landscapes of Recombination in Drosophila melanogaster. Plos Genetics 8.

Danecek P, Auton A, Abecasis G, Albers CA, Banks E, DePristo MA, Handsaker RE, Lunter G, Marth GT, Sherry ST, et al. 2011. The variant call format and VCFtools. Bioinformatics 27:2156–2158.

Diaz-Gonzalez J, Vazquez JF, Albornoz J, Dominguez A. 2011. Long-term evolution of the roo transposable element copy number in mutation accumulation lines of Drosophila melanogaster. Genetics Research 93:181–187.

Ewing AD. 2015. Transposable element detection from whole genome sequence data. Mobile DNA 6.

Gao ZY, Moorjani P, Sasani TA, Pedersen BS, Quinlan AR, Jorde LB, Amster G, Przeworski M. 2019. Overlooked roles of DNA damage and maternal age in generating human germline mutations. Proceedings of the National Academy of Sciences of the United States of America 116:9491–9500.

Guerreiro MPG. 2012. What makes transposable elements move in the Drosophila genome? Heredity 108:461–468.

Guerreiro MPG, Chavez-Sandoval BE, Balanya J, Serra L, Fontdevila A. 2008. Distribution of the transposable elements bilbo and gypsy in original and colonizing populations of Drosophila subobscura. Bmc Evolutionary Biology 8.

Guerreiro MPG, Fontdevila A. 2011. Osvaldo and Isis retrotransposons as markers of the Drosophila buzzatii colonisation in Australia. Bmc Evolutionary Biology 11.

Haag-Liautard C, Dorris M, Maside X, Macaskill S, Halligan DL, Charlesworth B, Keightley PD. 2007. Direct estimation of per nucleotide and genomic deleterious mutation rates in Drosophila. Nature 445:82–85.

Hadfield J. 2014. MCMCglmm course notes. available at: cran.r-project.org/web/packages/MCMCglmm/vignettes/CourseNotes.pdf.

Hadfield JD. 2010. MCMC Methods for Multi-Response Generalized Linear Mixed Models: The MCMCglmm R Package. Journal of Statistical Software 33:1–22.

Heil CSS, Ellison C, Dubin M, Noor MAF. 2015. Recombining without Hotspots: A Comprehensive Evolutionary Portrait of Recombination in Two Closely Related Species of Drosophila. Genome biology and evolution 7:2829–2842.

Hemmer LW, Dias GB, Smith B, Van Vaerenberghe K, Howard A, Bergman CM, Blumenstiel JP. 2020. Hybrid dysgenesis in Drosophila virilis results in clusters of mitotic recombination and loss-of-heterozygosity but leaves meiotic recombination unaltered. Mobile DNA 11.

Hirsch CD, Springer NM. 2017. Transposable element influences on gene expression in plants. Biochim Biophys Acta Gene Regul Mech 1860:157–165.

Horvath V, Merenciano M, Gonzalez J. 2017. Revisiting the Relationship between Transposable Elements and the Eukaryotic Stress Response. Trends in Genetics 33:832–841.

Huang W, Lyman RF, Lyman RA, Carbone MA, Harbison ST, Magwire MM, Mackay TFC. 2016. Spontaneous mutations and the origin and maintenance of quantitative genetic variation. Elife 5.

Hunter CM, Huang W, Mackay TFC, Singh ND. 2016. The Genetic Architecture of Natural Variation in Recombination Rate in Drosophila melanogaster. Plos Genetics 12.

Hunter CM, Singh ND. 2014. Do Males Matter? Testing the Effects of Male Genetic Background on Female Meiotic Crossover Rates in Drosophila Melanogaster. Evolution 68:2718–2726.

Jonsson H, Sulem P, Kehr B, Kristmundsdottir S, Zink F, Hjartarson E, Hardarson MT, Hjorleifsson KE, Eggertsson HP, Gudjonsson SA, et al. 2017. Parental influence on human germline de novo mutations in 1,548 trios from Iceland. Nature 549:519–522.

Kapitonov VV, Jurka J. 2003. Molecular paleontology of transposable elements in the Drosophila melanogaster genome. Proceedings of the National Academy of Sciences of the United States of America 100:6569–6574.

Kaplanis J, Ide B, Sanghvi R, Neville M, Danecek P, Coorens T, Prigmore E, Short P, Gallone G, McRae J, et al. 2021. Genetic and chemotherapeutic causes of germline hypermutation. bioRxiv:2021.2006.2001.446180.

Kapopoulou A, Pfeifer SP, Jensen JD, Laurent S. 2018. The Demographic History of African Drosophila melanogaster. Genome Biol Evol 10:2338–2342.

Kapun M, van Schalkwyk H, McAllister B, Flatt T, Schlotterer C. 2014. Inference of chromosomal inversion dynamics from Pool-Seq data in natural and laboratory populations of Drosophila melanogaster. Molecular Ecology 23:1813–1827.

Keightley PD, Ness RW, Halligan DL, Haddrill PR. 2014. Estimation of the Spontaneous Mutation Rate per Nucleotide Site in a Drosophila melanogaster Full-Sib Family. Genetics 196:313–320.

Keightley PD, Pinharanda A, Ness RW, Simpson F, Dasmahapatra KK, Mallet J, Davey JW, Jiggins CD. 2015. Estimation of the spontaneous mutation rate in Heliconius melpomene. Molecular Biology and Evolution 32:239–243.

Keightley PD, Trivedi U, Thomson M, Oliver F, Kumar S, Blaxter ML. 2009. Analysis of the genome sequences of three Drosophila melanogaster spontaneous mutation accumulation lines. Genome Research 19:1195–1201.

Kimura M. 1991. The Neutral Theory of Molecular Evolution - a Review of Recent-Evidence. Japanese Journal of Genetics 66:367–386.

Kong A, Frigge ML, Masson G, Besenbacher S, Sulem P, Magnusson G, Gudjonsson SA, Sigurdsson A, Jonasdottir A, Jonasdottir A, et al. 2012. Rate of de novo mutations and the importance of father’s age to disease risk. Nature 488:471–475.

Krasovec M. 2021. The spontaneous mutation rate of Drosophila pseudoobscura. G3-Genes Genomes Genetics 11.

Li H. 2013. Aligning sequence reads, clone sequences and assembly contigs with BWA-MEM. arXiv preprint arXiv:1303.3997.

Li H. 2011. A statistical framework for SNP calling, mutation discovery, association mapping and population genetical parameter estimation from sequencing data. Bioinformatics 27:2987–2993.

Lindsay SJ, Rahbari R, Kaplanis J, Keane T, Hurles ME. 2019. Similarities and differences in patterns of germline mutation between mice and humans. Nature Communications 10.

Liu H, Jia Y, Sun X, Tian D, Hurst LD, Yang S. 2017. Direct Determination of the Mutation Rate in the Bumblebee Reveals Evidence for Weak Recombination-Associated Mutation and an Approximate Rate Constancy in Insects. Molecular Biology and Evolution 34:119–130.

Lynch M. 2010. Evolution of the mutation rate. Trends in Genetics 26:345–352.

Lynch M, Ackerman MS, Gout JF, Long H, Sung W, Thomas WK, Foster PL. 2016. Genetic drift, selection and the evolution of the mutation rate. Nature Reviews Genetics 17:704–714.

Martin M. 2011. Cutadapt removes adapter sequences from high-throughput sequencing reads. EMBnet. journal 17:10–12.

Maside X, Assimacopoulos S, Charlesworth B. 2000. Rates of movement of transposable elements on the second chromosome of Drosophila melanogaster. Genetical Research 75:275–284.

McKenna A, Hanna M, Banks E, Sivachenko A, Cibulskis K, Kernytsky A, Garimella K, Altshuler D, Gabriel S, Daly M, et al. 2010. The Genome Analysis Toolkit: a MapReduce framework for analyzing next-generation DNA sequencing data. Genome Research 20:1297–1303.

Merel V, Boulesteix M, Fablet M, Vieira C. 2020. Transposable elements in Drosophila. Mob DNA 11:23.

Miles A, Harding NJ, Botta G, Clarkson CS, Antao T, Kozak K, Schrider DR, Kern AD, Redmond S, Sharakhov I, et al. 2017. Genetic diversity of the African malaria vector Anopheles gambiae. Nature 552:96–+.

Milholland B, Dong X, Zhang L, Hao XX, Suh Y, Vijg J. 2017. Differences between germline and somatic mutation rates in humans and mice. Nature Communications 8.

Miller DE, Smith CB, Kazemi NY, Cockrell AJ, Arvanitakas AV, Blumenstiel JP, Jaspersen SL, Hawley RS. 2016. Whole-Genome Analysis of Individual Meiotic Events in Drosophila melanogaster Reveals That Noncrossover Gene Conversions Are Insensitive to Interference and the Centromere Effect. Genetics 203:159–171.

Misof B, Liu S, Meusemann K, Peters RS, Donath A, Mayer C, Frandsen PB, Ware J, Flouri T, Beutel RG, et al. 2014. Phylogenomics resolves the timing and pattern of insect evolution. Science 346:763–767.

Morgan TH. 1912. Complete linkage in the second chromosome of the male of Drosophila. Science 36:719–720.

Ness RW, Morgan AD, Vasanthakrishnan RB, Colegrave N, Keightley PD. 2015. Extensive de novo mutation rate variation between individuals and across the genome of Chlamydomonas reinhardtii. Genome Research 25:1739–1749.

Nuzhdin SV, Mackay TFC. 1995. The Genomic Rate of Transposable Element Movement in Drosophila-Melanogaster. Molecular Biology and Evolution 12:180–181.

O’Roak BJ, Vives L, Girirajan S, Karakoc E, Krumm N, Coe BP, Levy R, Ko A, Lee C, Smith JD, et al. 2012. Sporadic autism exomes reveal a highly interconnected protein network of de novo mutations. Nature 485:246–U136.

Obbard DJ, Maclennan J, Kim KW, Rambaut A, O’Grady PM, Jiggins FM. 2012. Estimating Divergence Dates and Substitution Rates in the Drosophila Phylogeny. Molecular Biology and Evolution 29:3459–3473.

Oppold AM, Pfenninger M. 2017. Direct estimation of the spontaneous mutation rate by short-term mutation accumulation lines in Chironomus riparius. Evolution Letters 1:86–92.

Parsch J, Novozhilov S, Saminadin-Peter SS, Wong KM, Andolfatto P. 2010. On the Utility of Short Intron Sequences as a Reference for the Detection of Positive and Negative Selection in Drosophila. Molecular Biology and Evolution 27:1226–1234.

Pasyukova EG, Nuzhdin SV, Filatov DA. 1998. The relationship between the rate of transposition and transposable element copy number for copia and Doc retrotransposons of Drosophila melanogaster. Genetics Research 72:1–11.

Pfeifer GP, You YH, Besaratinia A. 2005. Mutations induced by ultraviolet light. Mutation Research-Fundamental and Molecular Mechanisms of Mutagenesis 571:19–31.

Rahbari R, Wuster A, Lindsay SJ, Hardwick RJ, Alexandrov LB, Turki SA, Dominiczak A, Morris A, Porteous D, Smith B, et al. 2016. Timing, rates and spectra of human germline mutation. Nat Genet 48:126–133.

Rech GE. 2021. Fasta sequences for the Drosophila melanogaster Manually Curated Transposable Elements (MCTE) library. In: DIGITAL.CSIC.

Redfield H. 1966. Delayed mating and the relationship of recombination to maternal age in Drosophila melanogaster. Genetics 53:593–607.

Rimmer A, Phan H, Mathieson I, Iqbal Z, Twigg SRF, Wilkie AOM, McVean G, Lunter G, Consortium W. 2014. Integrating mapping-, assembly-and haplotype-based approaches for calling variants in clinical sequencing applications. Nature Genetics 46:912–918.

Robinson JT, Thorvaldsdottir H, Winckler W, Guttman M, Lander ES, Getz G, Mesirov JP. 2011. Integrative genomics viewer. Nature Biotechnology 29:24–26.

Roles AJ, Conner JK. 2008. Fitness effects of mutation accumulation in a natural outbred population of wild radish (raphanus raphanistrum): Comparison of field and greenhouse environments. Evolution 62:1066–1075.

Schrider DR, Houle D, Lynch M, Hahn MW. 2013. Rates and Genomic Consequences of Spontaneous Mutational Events in Drosophila melanogaster. Genetics 194:937–954.

Schwarz F, Wierzbicki F, Senti KA, Kofler R. 2021. Tirant Stealthily Invaded Natural Drosophila melanogaster Populations during the Last Century. Molecular Biology and Evolution 38:1482–1497.

Segurel L, Wyman MJ, Przeworski M. 2014. Determinants of Mutation Rate Variation in the Human Germline. Annual Review of Genomics and Human Genetics, Vol 15 15:47–70.

Sharp NP, Agrawal AF. 2016. Low Genetic Quality Alters Key Dimensions of the Mutational Spectrum. Plos Biology 14.

Stapley J, Feulner PGD, Johnston SE, Santure AW, Smadja CM. 2017. Variation in recombination frequency and distribution across eukaryotes: patterns and processes. Philos Trans R Soc Lond B Biol Sci 372.

Stevison LS, Noor MAF. 2010. Genetic and Evolutionary Correlates of Fine-Scale Recombination Rate Variation in Drosophila persimilis. Journal of Molecular Evolution 71:332–345.

Stocker AJ, Rusuwa BB, Blacket MJ, Frentiu FD, Sullivan M, Foley BR, Beatson S, Hoffmann AA, Chenoweth SF. 2012. Physical and Linkage Maps for Drosophila serrata, a Model Species for Studies of Clinal Adaptation and Sexual Selection. G3 (Bethesda) 2:287–297.

Sung W, Ackerman MS, Dillon MM, Platt TG, Fuqua C, Cooper VS, Lynch M. 2016. Evolution of the Insertion-Deletion Mutation Rate Across the Tree of Life. G3-Genes Genomes Genetics 6:2583–2591.

Sung W, Ackerman MS, Gout JF, Miller SF, Williams E, Foster PL, Lynch M. 2015. Asymmetric Context-Dependent Mutation Patterns Revealed through Mutation-Accumulation Experiments. Molecular Biology and Evolution 32:1672–1683.

Suvorov A, Kim BY, Wang J, Armstrong EE, Peede D, D’Agostino ERR, Price DK, Waddell P, Lang M, Courtier-Orgogozo V, et al. 2022. Widespread introgression across a phylogeny of 155 Drosophila genomes. Current Biology 32:111–123 e115.

Taylor F, Tyekucheva S, Zody M, Chiaromonte F, Makova KD. 2006. Strong and weak male mutation bias at different sites in the primate genomes: Insights from the human-chimpanzee comparison. Molecular Biology and Evolution 23:565–573.

Thomas GWC, Wang RJ, Puri A, Harris RA, Raveendran M, Hughes DST, Murali SC, Williams LE, Doddapaneni H, Muzny DM, et al. 2018. Reproductive Longevity Predicts Mutation Rates in Primates. Current Biology 28:3193–3197.

True JR, Mercer JM, Laurie CC. 1996. Differences in crossover frequency and distribution among three sibling species of Drosophila. Genetics 142:507–523.

Uchimura A, Higuchi M, Minakuchi Y, Ohno M, Toyoda A, Fujiyama A, Miura I, Wakana S, Nishino J, Yagi T. 2015. Germline mutation rates and the long-term phenotypic effects of mutation accumulation in wild-type laboratory mice and mutator mice. Genome Research 25:1125–1134.

Waldvogel AM, Pfenninger M. 2021. Temperature dependence of spontaneous mutation rates. Genome Research 31:1582–1589.

Webster MT, Hurst LD. 2012. Direct and indirect consequences of meiotic recombination: implications for genome evolution. Trends in Genetics 28:101–109.

Wells JN, Feschotte C. 2020. A Field Guide to Eukaryotic Transposable Elements. Annu Rev Genet 54:539–561.

Yang JA, Lee SH, Goddard ME, Visscher PM. 2011. GCTA: A Tool for Genome-wide Complex Trait Analysis. American Journal of Human Genetics 88:76–82.

Yang S, Wang L, Huang J, Zhang X, Yuan Y, Chen JQ, Hurst LD, Tian D. 2015. Parent-progeny sequencing indicates higher mutation rates in heterozygotes. Nature 523:463–467.

Yoder AD, Tiley GP. 2021. The challenge and promise of estimating the de novo mutation rate from whole-genome comparisons among closely related individuals. Molecular Ecology 30:6087–6100.

