## Supplementary figures and images for "Variation in mutation, recombination, and transposition rates in *Drosophila melanogaster* and *Drosophila simulans*"

### supplementary fig. S1

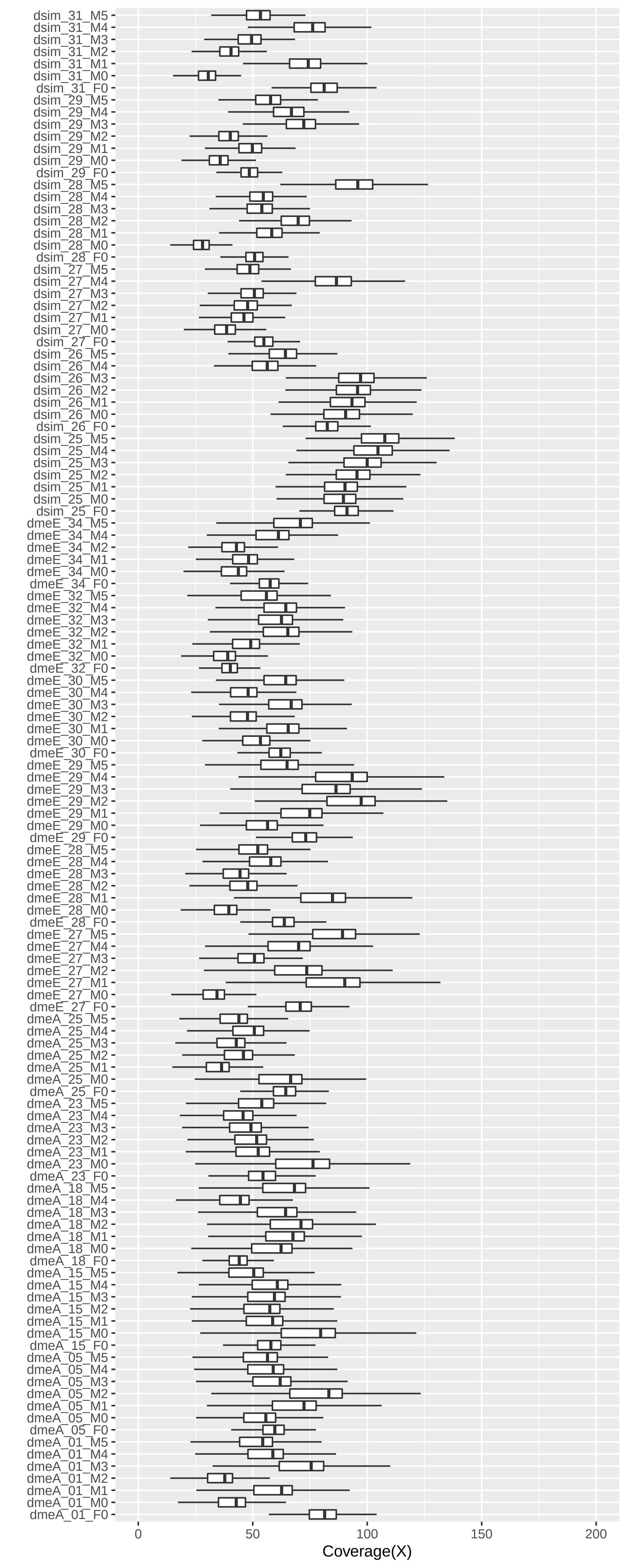

### supplementary fig. S2

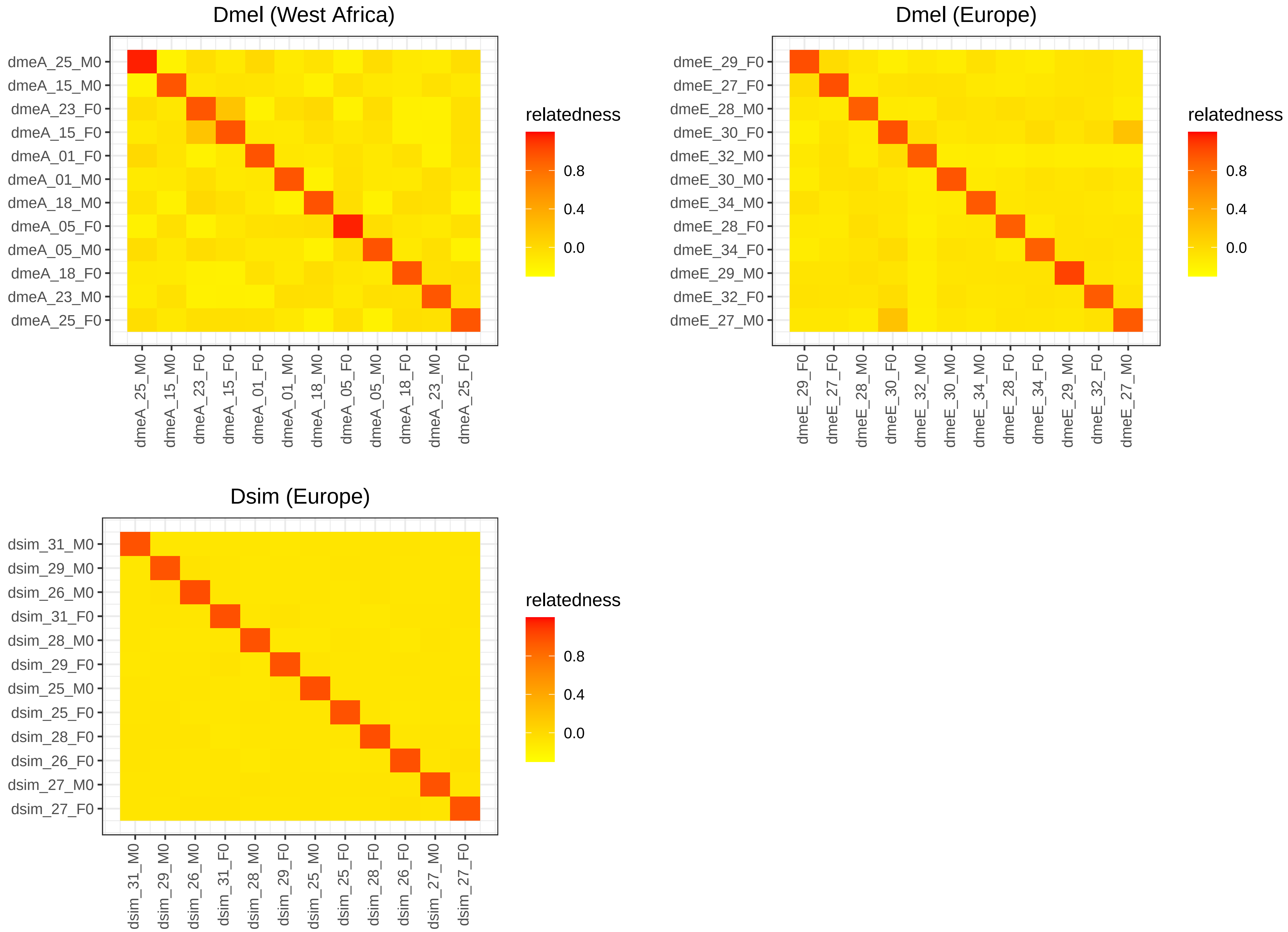

### supplementary fig. S3

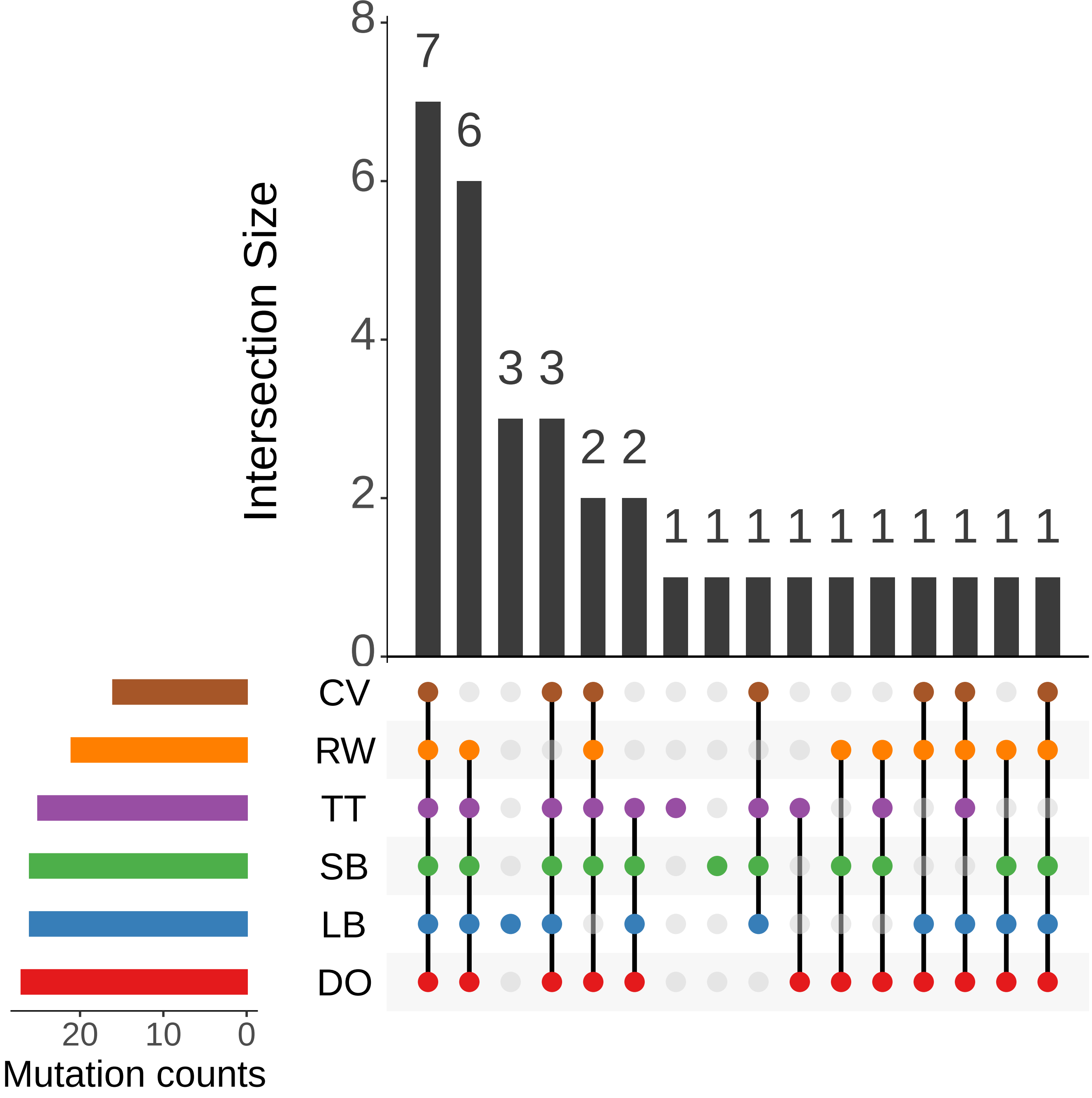

### supplementary fig. S4

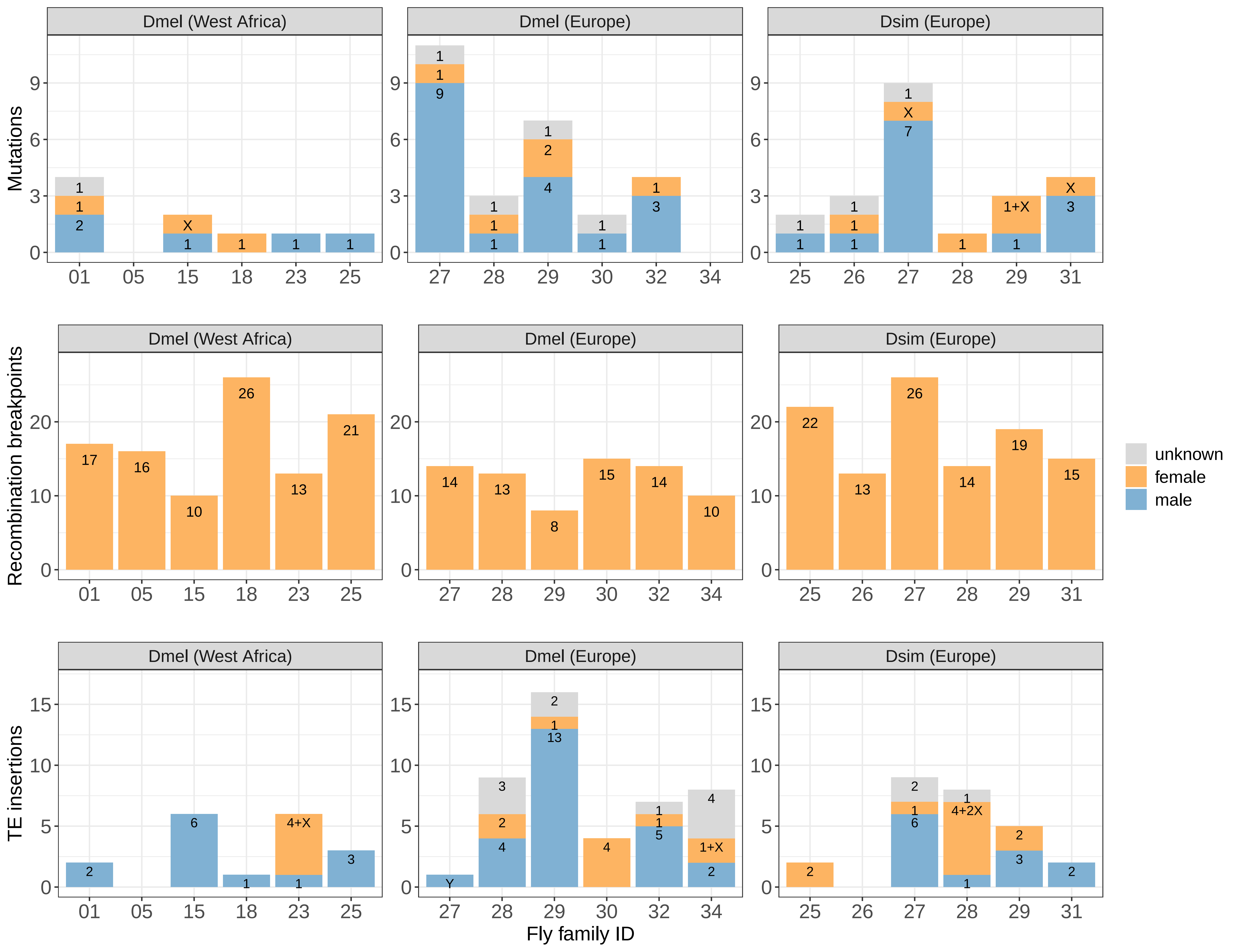

### supplementary fig. S5

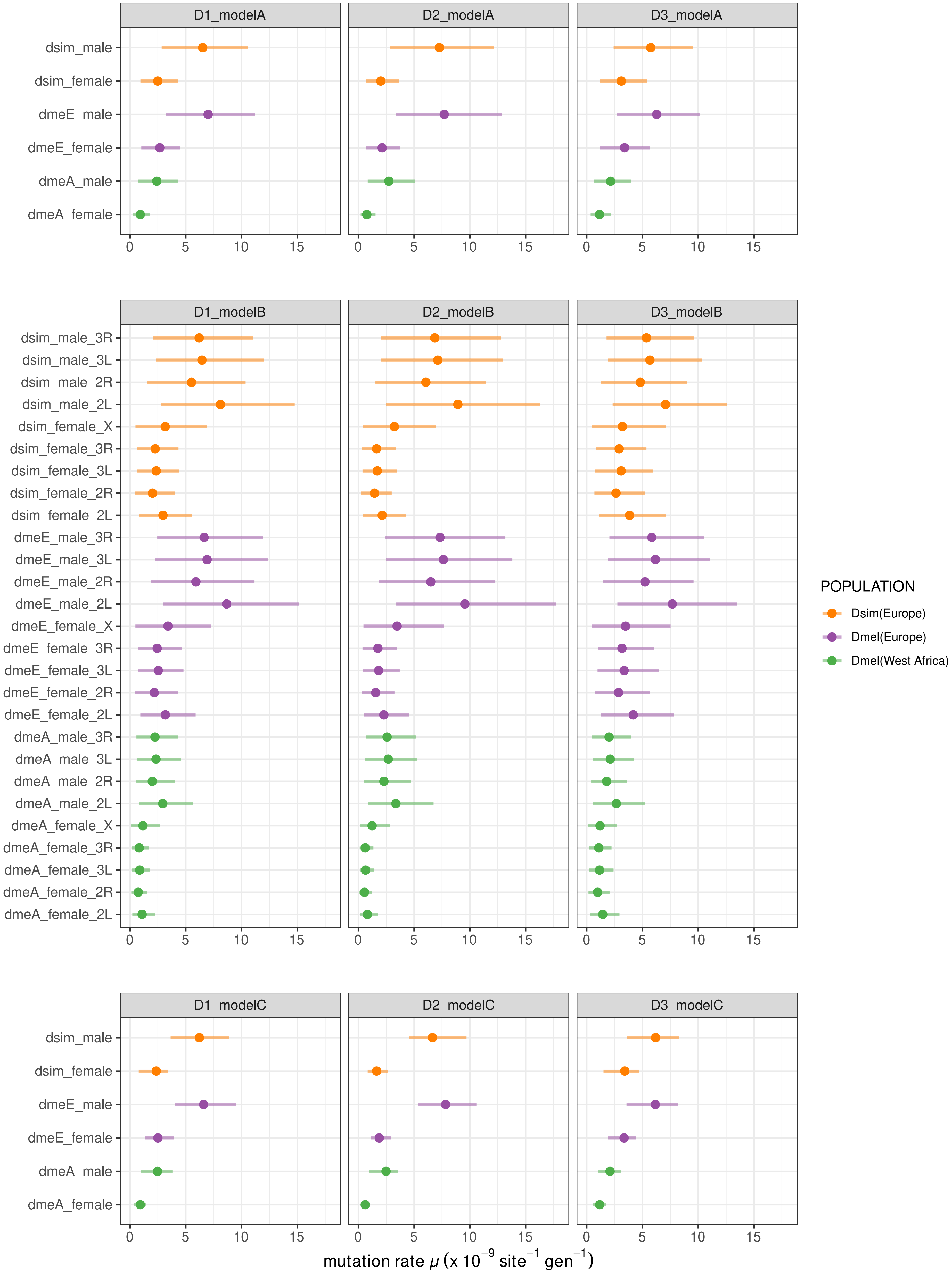

### supplementary fig. S7

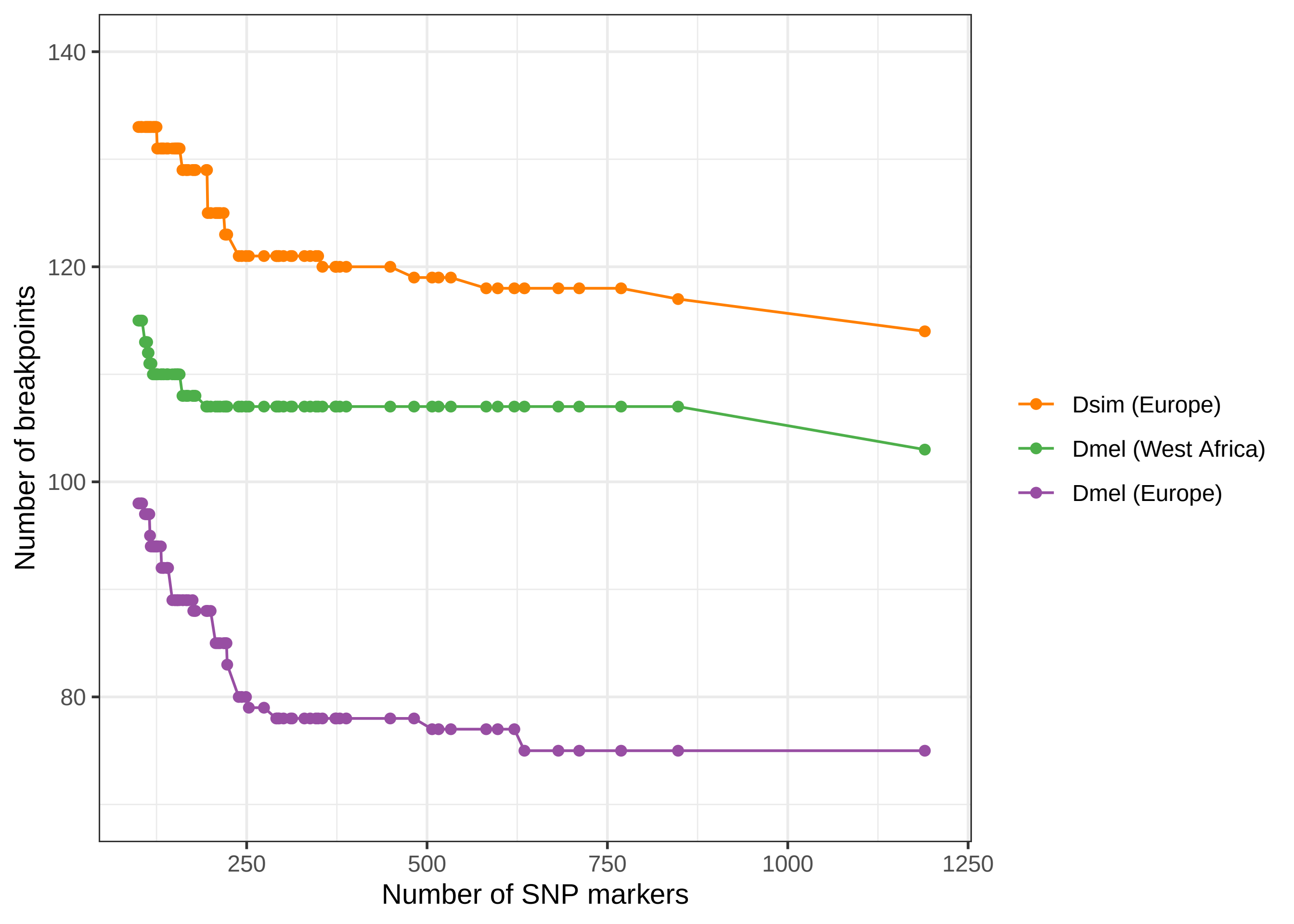

### supplementary fig. S8

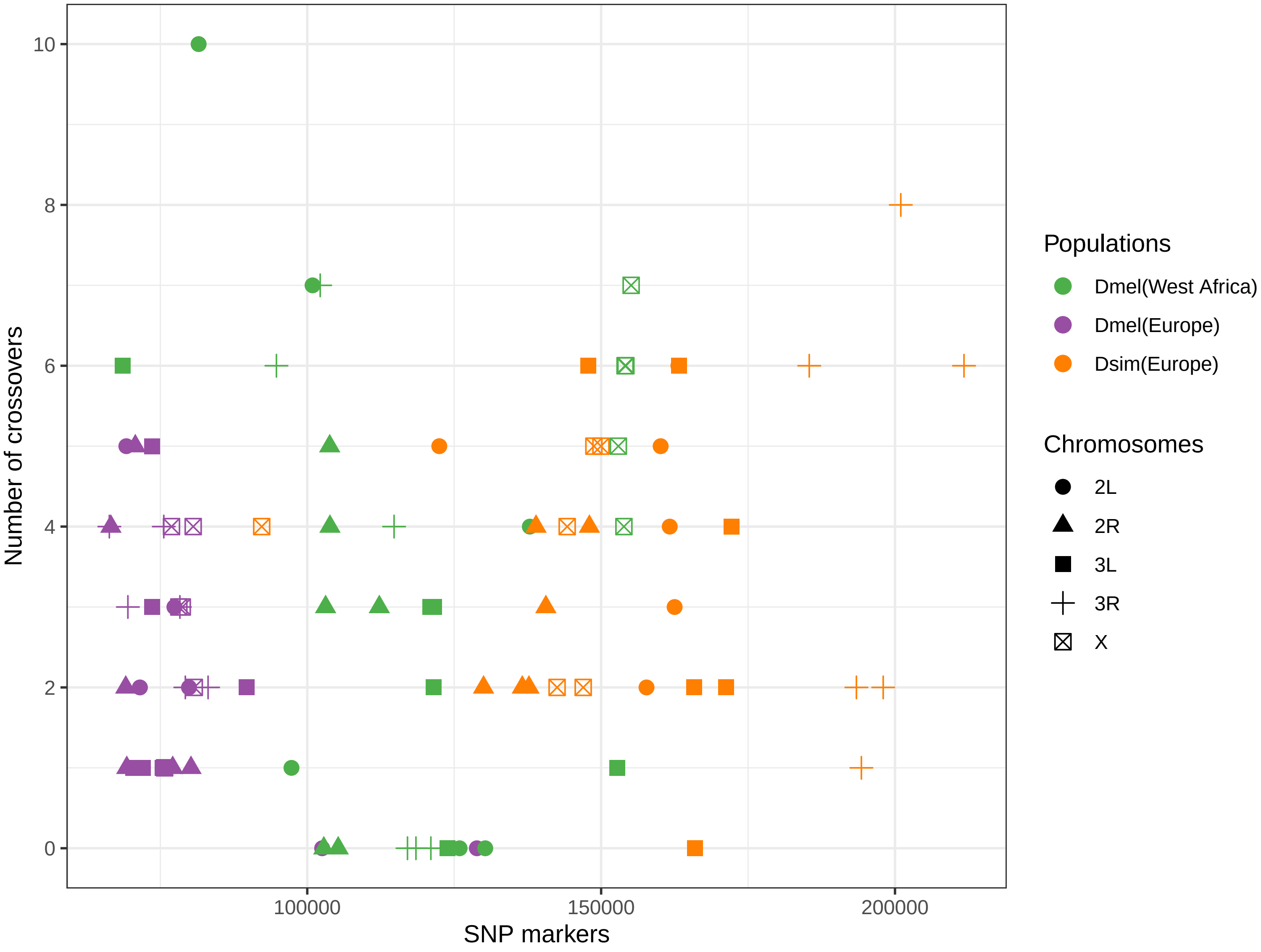

### supplementary fig. S9

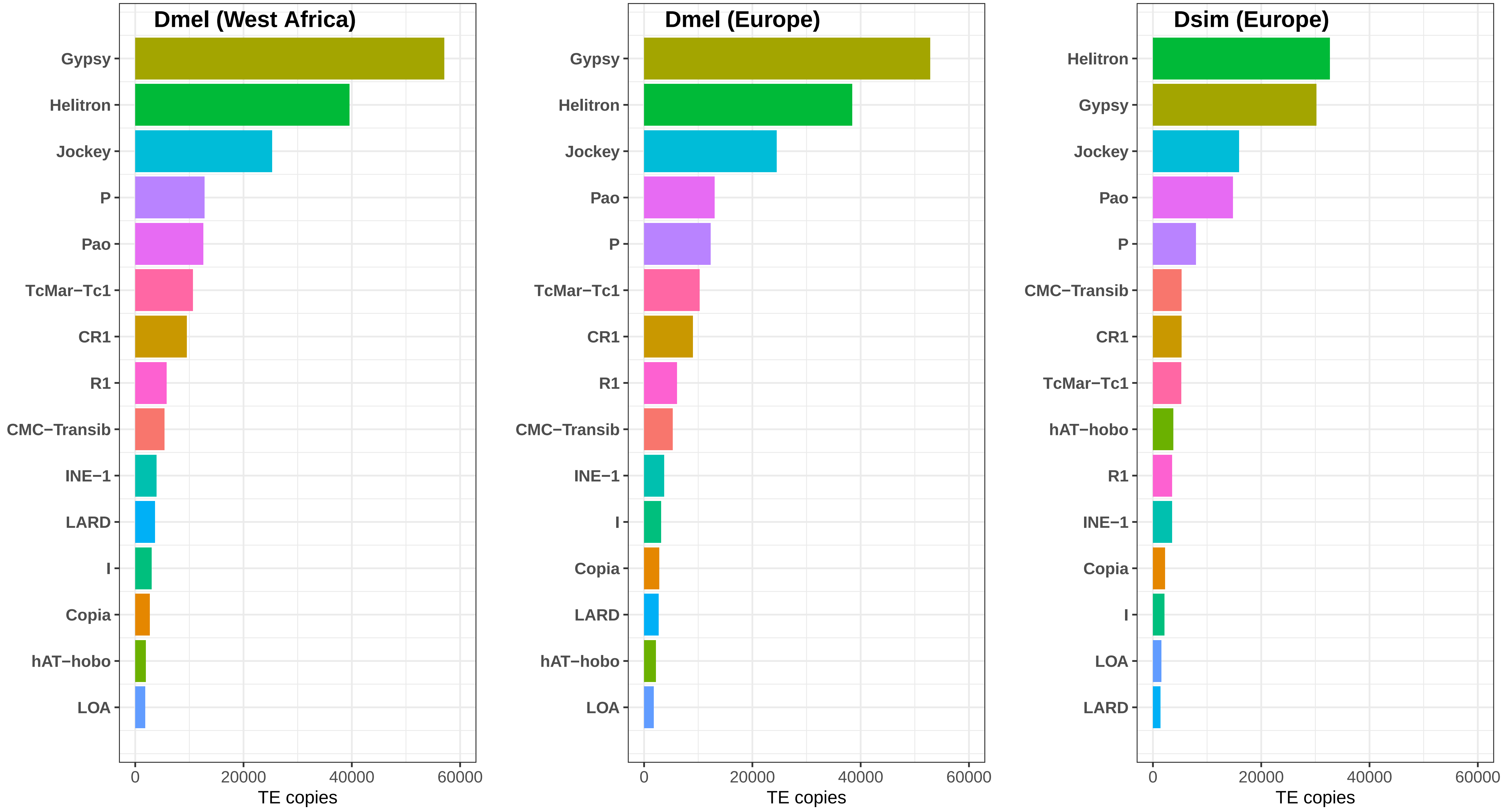

### supplementary fig. S10

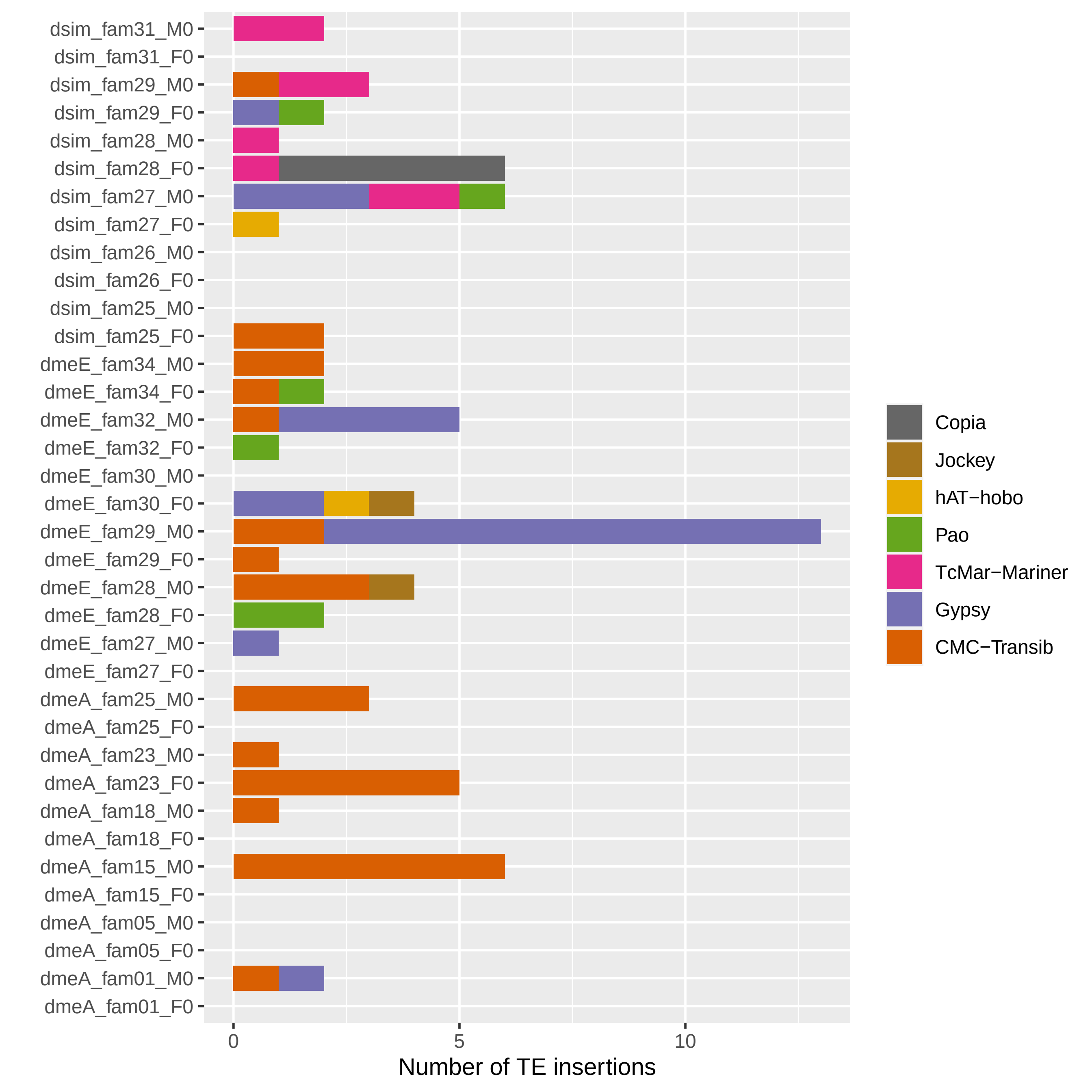
