## supplementary fig. S6 for "Variation in mutation, recombination, and transposition rates in *Drosophila melanogaster* and *Drosophila simulans*"

*D. melanogaster* (West Africa)

2L	2R	3L	3R	X

2L	2R	3L	3R

Fam25	Fam23	Fam18	Fam15	Fam05	Fam01

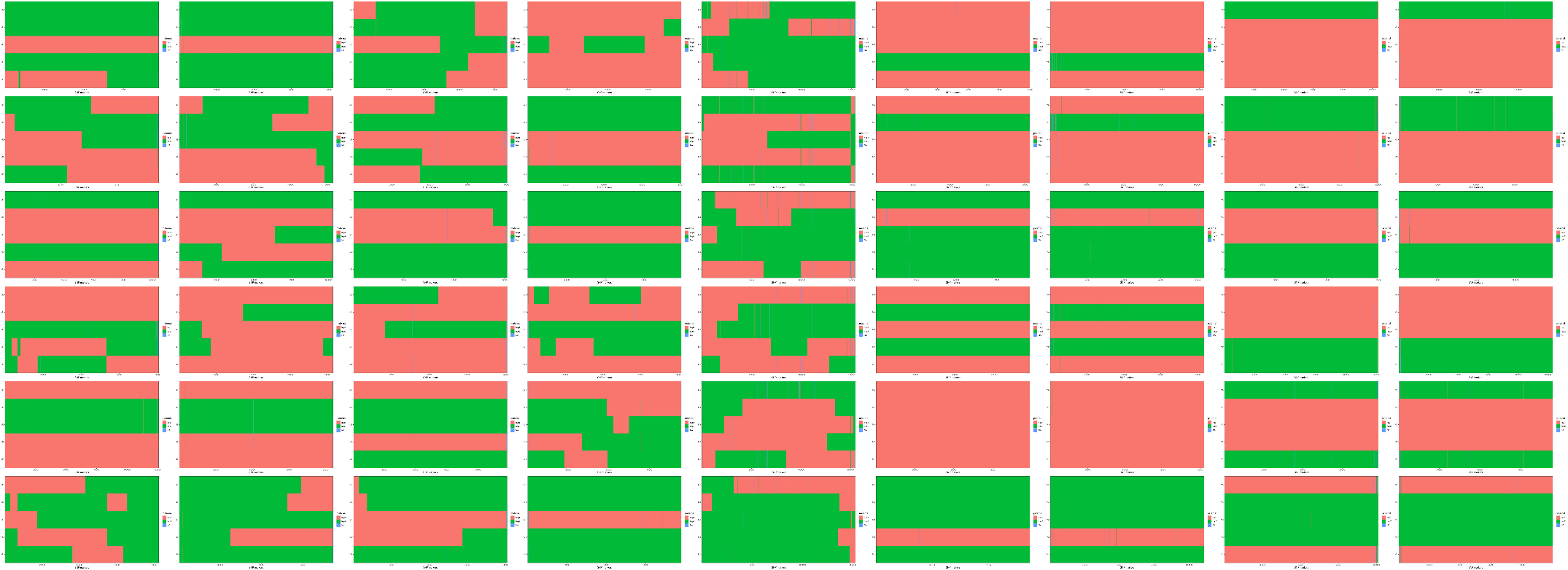

paternal

maternal

*D. melanogaster* (Europe)

2L	2R	3L	3R

2L	2R	3L	3R	X

Fam34	Fam32	Fam30	Fam29	Fam28	Fam27

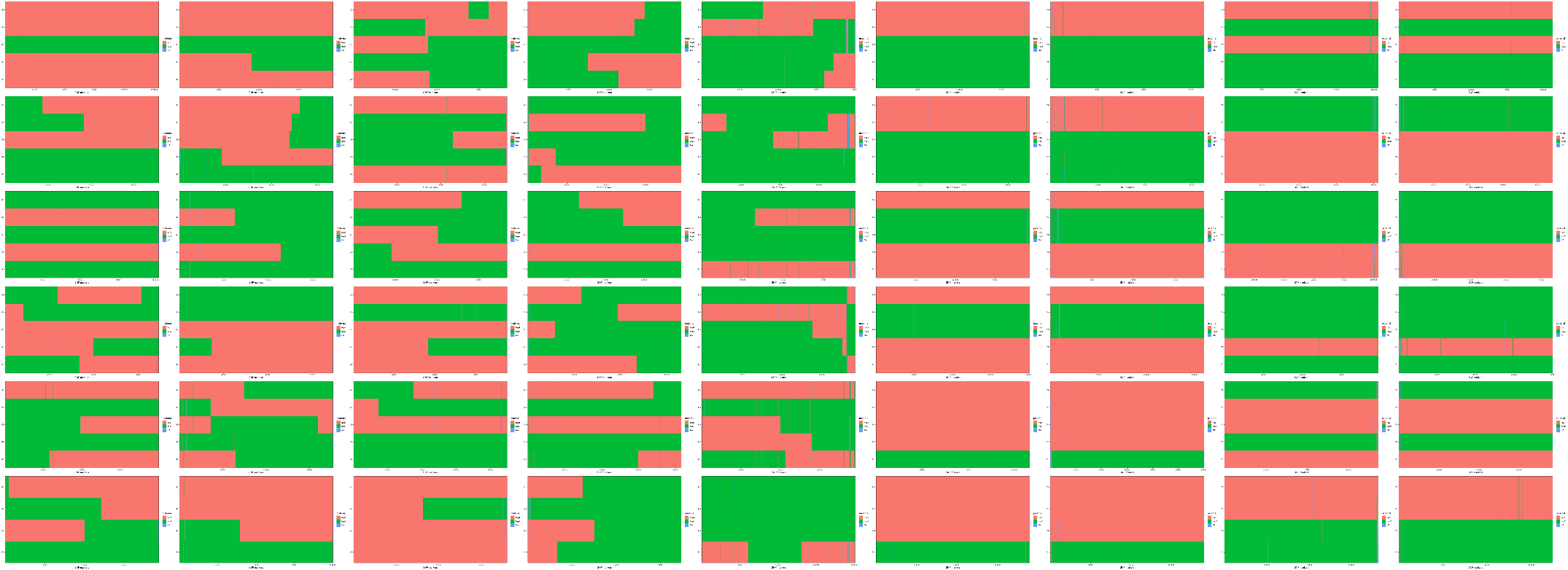

paternal

maternal

*D. simulans* (Europe)

2L	2R	3L	3R

2L	2R	3L	3R	X

Fam31	Fam29	Fam28	Fam27	Fam26	Fam25

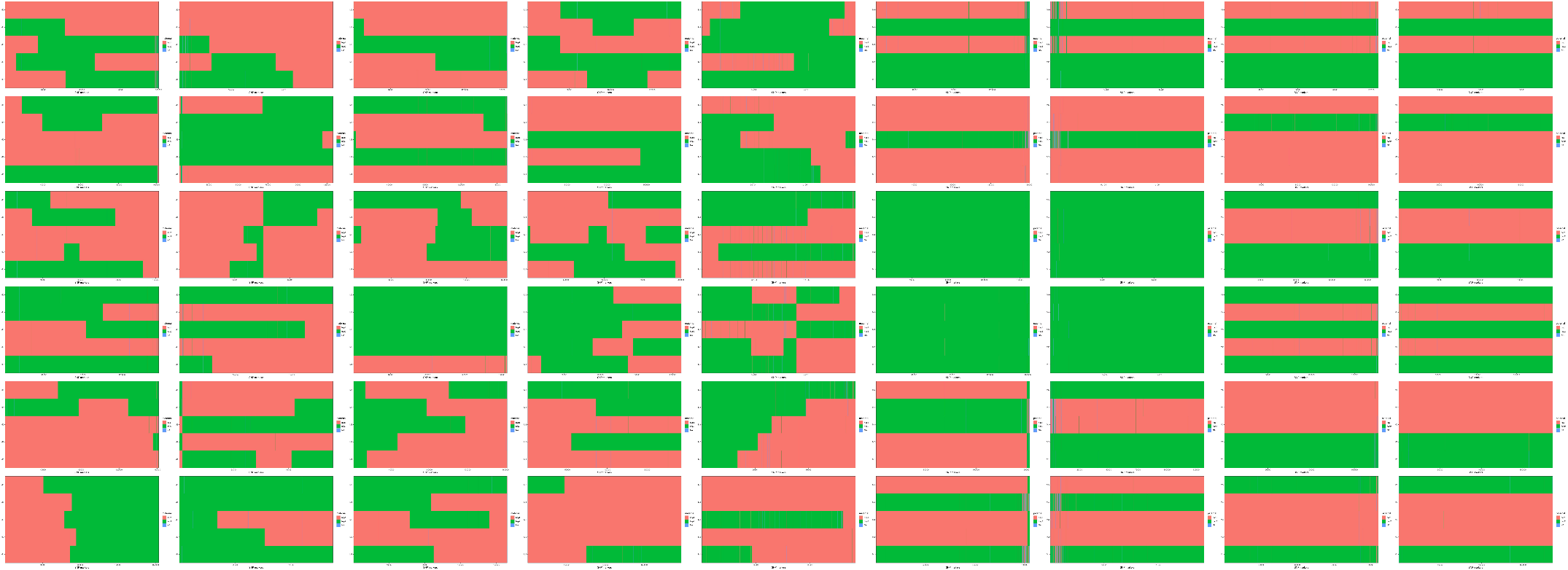

paternal

maternal
